# scTriangulate, a game-theory based framework for optimal solutions of uni- and multimodal single-cell data

**DOI:** 10.1101/2021.10.16.464640

**Authors:** Guangyuan Li, Baobao Song, Harinder Singh, V. B. Surya Prasath, H. Leighton Grimes, Nathan Salomonis

**Affiliations:** Division of Biomedical Informatics, Cincinnati Children’s Hospital Medical Center, Cincinnati, OH, USA; Department of Biomedical Informatics, College of Medicine, University of Cincinnati, OH, 45267 USA; Center for Systems Immunology, Departments of Immunology and Computational and Systems Biology, University of Pittsburgh, Pittsburgh, PA, USA; Division of Immunobiology, Cincinnati Children’s Hospital Medical Center, Cincinnati, OH, USA; Department of Pediatrics, University of Cincinnati School of Medicine, Cincinnati, Ohio, USA; Department of Electrical Engineering and Computer Science, University of Cincinnati, OH 45221 USA

## Abstract

Decisively delineating cell identities from uni- and multimodal single-cell datasets is complicated by diverse modalities, clustering methods, and reference atlases. We describe scTriangulate, a cooperative game-theory framework that mixes-and-matches multiple clustering results, modalities, associated algorithms, and resolutions to achieve an optimal solution. Rather than ensemble approaches which select the “consensus”, scTriangulate picks the most stable solution. When evaluated on diverse multimodal technologies, scTriangulate outperforms alternative approaches to identify consistent high-confidence novel cell populations and modality-specific subtypes. Unlike existing integration strategies that rely on modality-specific joint embedding or geometric graphs, scTriangulate makes no assumption about the distributions of raw underlying values. As a result, this approach can solve unprecedented integration challenges, including the ability to automate reference cell-atlas construction, resolve clonal architecture within molecularly defined cell-populations and subdivide clusters to discover novel splicing-defined disease subtypes. scTriangulate is a flexible strategy for unified integration of single-cell or multimodal clustering solutions, from nearly unlimited sources.

## INTRODUCTION

Single-cell genomics has significantly evolved from its original introduction, enabling the analysis of millions of cells and many molecular modalities in complex organisms. The repertoire of simultaneous detectable modalities in a single-cell continues to grow, which now include RNA abundance, DNA-accessibility, surface antigen expression, somatic mutations and epigenetic modifications among others ^1–4^. To identify cell populations defined from multiple modalities in a single experiment, new bioinformatics approaches have begun to emerge. These include methods such as Seurat-WNN, which applies a weight-nearest neighbor (WNN) approach to integrate clustering results from two or more supported modalities ^5^, and TotalVI which leverages a variational autoencoder to infer the joint latent space considering RNA and surface protein expression ^6^. Notably, such methods can produce improved results over single-modality clustering methods and new gene regulatory insights. However, these and other methods are: 1) limited to specific modalities, 2) cannot jointly consider prior annotations from label transfer methods and 3) are limited to specific types of unsupervised clustering algorithms for all modalities. Importantly, such existing integration methods inherently produce extremely variable results, dependent on the software resolution selected ^7–9^.

It is likely that no single-resolution in multimodal single-cell analyses will identify the most discrete cell-states in different modalities, but rather different resolutions will highlight informative populations (i.e., sub-clusters, clonal specific impacts) when considered for a range of possible resolutions. These problems are compounded with the publication of hundreds of single-cell atlases which pose the enormous challenge of combining and reconciling population labels from prior predictions (unimodal or multimodal), while including those predicted from parallel unsupervised analyses. While consensus clustering through ensemble approaches, such as the Meta-CLustering Algorithm (MCLA), provide a possible solution to this problem, these methods only consider agreement between diverse clustering results or modalities, without considering statistical measures of cluster stability or the specificity of markers in individual populations^10^. Such stability metrics, including new robust machine-learning approaches, such as the Single Cell Clustering Assessment Framework (SCCAF) algorithm ^11^, are an important advance to resolve this challenge. Hence, to address these issues, there is a significant need for integration approaches that: a) are agnostic to the modality/algorithm/resolution considered, b) integrate between different clustering solutions (mix-and-match) and c) consider the biological stability of each cluster relative to all possible considered alternatives.

Here we describe scTriangulate, a new conceptual framework and associated computational workflow that overcomes these challenges by using game theory in combination with recent methods to compute cluster-stability. While game theory has been leveraged extensively in multiple domains, including economics, military strategy, transportation, and genetics, it has not been previously exploited in single-cell genomics ^12,13^. scTriangulate systematically mixes-and-matches cellular clusters generated by different algorithms spanning different modalities and resolutions to arrive at optimal clustering solutions (based upon cluster stability) (**Figure 1**). We consider each set of cluster annotations as an individual player and leverage the Shapley value from cooperative game theory to determine the relative importance of each set of cluster annotations^14^. scTriangulate considers distinct clusters produced from different unsupervised software clustering resolutions, clusters obtained from different single-cell modalities (e.g., RNA, ATAC), or annotations projected from well-defined reference cell atlases. While other multimodal methods integrate at the data-level (through either a low-dimensional latent space ^6^ or through a geometric graph ^5^), scTriangulate integrates results at a decision-level to reconcile conflicting cluster-label assignments (**Supplementary Figure 1**). scTriangulate is integrated seamlessly into the highly adopted scanpy python framework, enabling end-to-end analysis of multimodal and single-cell data. This flexible design makes it adaptable to new modalities (e.g., splicing, mutations), stability metrics, automated sub-clustering (e.g., sub-clonal analyses) and new types of research questions. Hence, this computational approach addresses numerous challenges not addressed by existing approaches, in particular: 1) overcoming arbitrary user-defined cluster resolutions that can result in biologically incoherent populations, 2) quantitative assessment of cluster reliability (stability, doublets) across the different modalities, 3) integration across algorithm or modality-specific results, and 4) ability to support new modalities. As it is not restricted to any specific algorithms, scTriangulate players can include existing multimodal clustering algorithms (e.g., Seurat-WNN, TotalVI), which may use their own specialized batch-effect correction methods. To enable careful evaluation of its output this tool builds a diversity of web-based interactive displays, to evaluate marker specificity, doublet predictions, cluster prediction confidence and the contribution of specific modalities to the final solutions.

**Figure 1.**
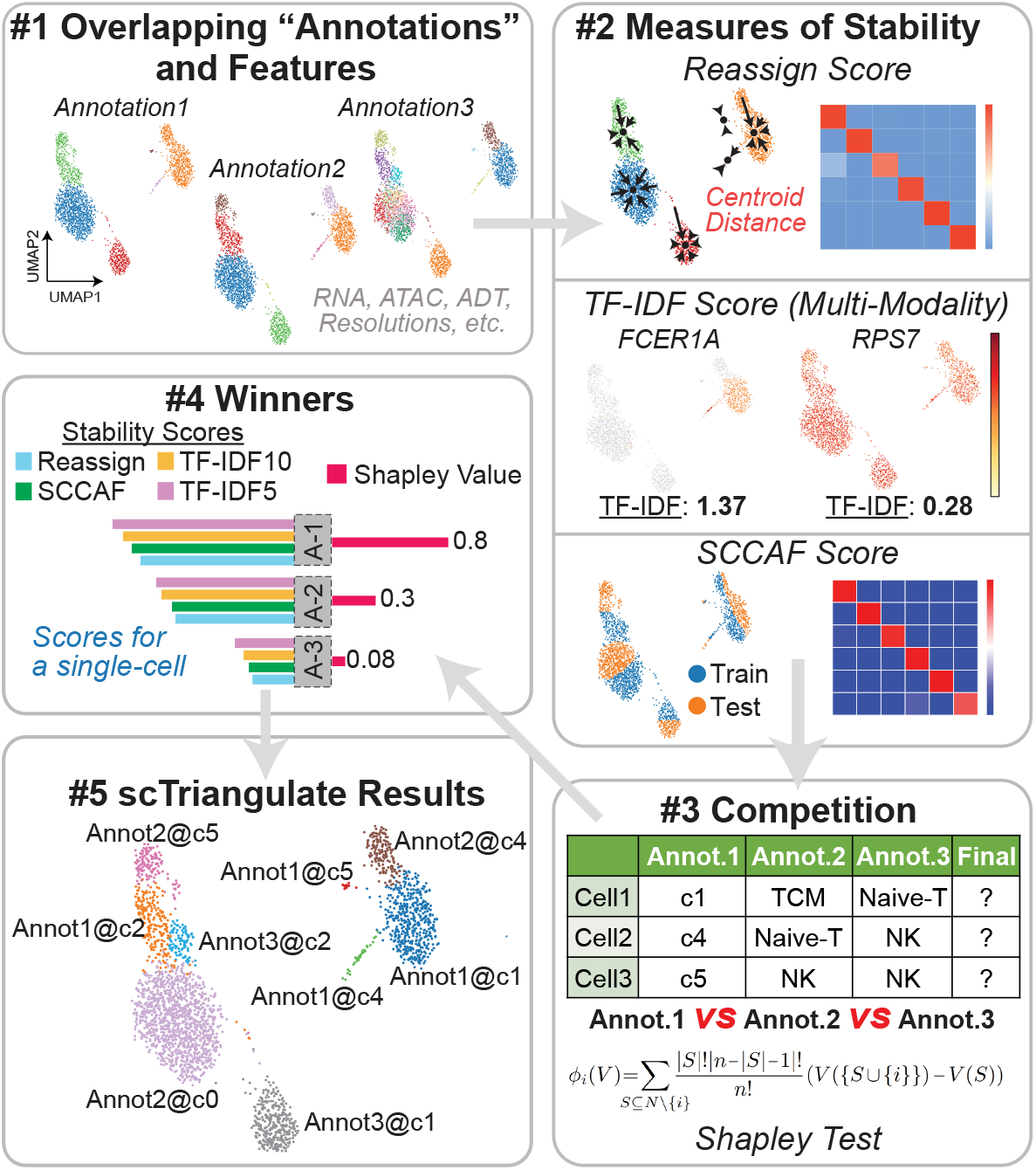
Resolving conflicting cluster identities from game-theory. Overview of scTriangulate. Using multiple competing annotation sources (algorithm, modality, resolution), different statistical stability metrics are computed for each cluster and cell. Each cell is assigned to the winning cluster label based on a computed Shapley Value (Steps 1-5, Methods). The reassign score considers the extent to which cells within a cluster can be reclassified to their own centroid based on nearest neighbor. The TF-IDF score corresponds to the statistical strength of the nth ranked feature (gene, ADT, peak) in a cluster. The SCCAF score implements a prior reliability metric (Single Cell Clustering Assessment Framework) that leverages multivariate multi-class logistic regression.

## RESULTS

### Uni- and Multimodal integration using game-theory

A fundamental assumption of scTriangulate is that no one algorithm or modality (set of molecular features) at a given resolution can provide optimal cell clustering solutions. Such stable solutions only emerge by combining (or triangulating) the low- and high-resolution cellular views derived from complementary approaches within a game-theoretic framework. At its core decision engine, scTriangulate resolves conflicting clusters for each single cell as a “game”, where each cluster label from a single annotation is a player. To ensure final clusters are robust, the stability of each cluster is assessed by an array of default and customizable stability metrics. Here, we assume that a valid cell population should possess a coherent molecular program (e.g., gene expression, epigenetic, post-transcriptional) to enable accurate reclassification, with distinct detected molecular markers. Specifically, we measure whether cells in a population can be correctly reclassified against its own centroid (reassign score) ^8^, the exclusivity of marker abundance (e.g., gene expression, epigenetic profile, post-transcriptional regulation) via term frequency-inverse document frequency (TF-IDF score) ^15^, and a combined multivariate single-cell clustering assessment framework (SCCAF score) ^11^. Diverse stability metrics were intentionally selected to assess stability from differential statistical perspectives. To assess stability across all modalities, all molecular features are combined into a single object and jointly evaluated.

While these stability ranks alone can be used to assess the validity of a cluster assignment for individual cells, the goal of scTriangulate is to assign each single cell the annotation with the highest collective rank for all stability metrics, while ensuring one algorithm does not always drive the decision. As inspiration, we considered the concept of inferred feature importance in machine learning, which is used in methods such as the SHAP value (SHapley Additive exPlanations) ^16^. The Shapley Value is used to gauge the relative contribution of each annotation to guarantee fair credit allocation ^14^. Rather than simply combine the stability ranks from each cell, Shapley maximizes overall cluster reliability by assessing the relative contribution of each player for all possible player combinations (coalitional iteration) for each cell. If a scTriangulate final cluster represents a small fraction of cells from the original cluster, the program automatically removes these cluster assignments (pruned, **Methods**) and re-classifies them against remaining final clusters. This filter ensures that outlier cells in a cluster, which do not selectively associate with a specific user provided annotation and are absorbed into neighboring clusters. For multi-modality datasets, features from each modality are jointly considered (e.g., RNA, ATAC, ADT, splicing, mutation), with the contribution of each modality to each final cluster reported (**Methods**). Hence, while an initial cluster input in scTriangulate may have been defined by one modality (e.g., RNA), features from other modalities can support the stability of that population (e.g., ATAC). scTriangulate has no upper limit on modalities because it selects/highlights biological features from individual modalities in obtaining the final clustering solution.

As an initial test of scTriangulate to identify the most valid cell populations from heterogeneous inputs, we created a simulated scRNA-Seq dataset using Splatter (**Figure 2a**). This simulated dataset was designed to include both broad cell-types and weekly distinguished cell states with unique gene expression as our ground-state truth. As the primary input to the software, we created alternative clustering solutions, in which each solution only partially matches our ground-state-truth (**Figure 2b**). Although the ground-state solution was not provided to scTriangulate, it was able to reproduce this solution by mixing and matching different clusters from the more broad (under-clustered) and granular (over-clustered) inputs, without including these artifacts (**Figure 2c**). Importantly, this outcome could not be obtained through tested conventional ensemble clustering techniques, which only consider consensus and not feature-based stability (**Figure 2d,e**). Additional simulation analyses demonstrate that scTriangulate can further integrate and resolve even more discrete gene expression differences by considering additional stability scores that promote weakly stable populations (**Supplementary Figure 2**).

**Figure 2.**
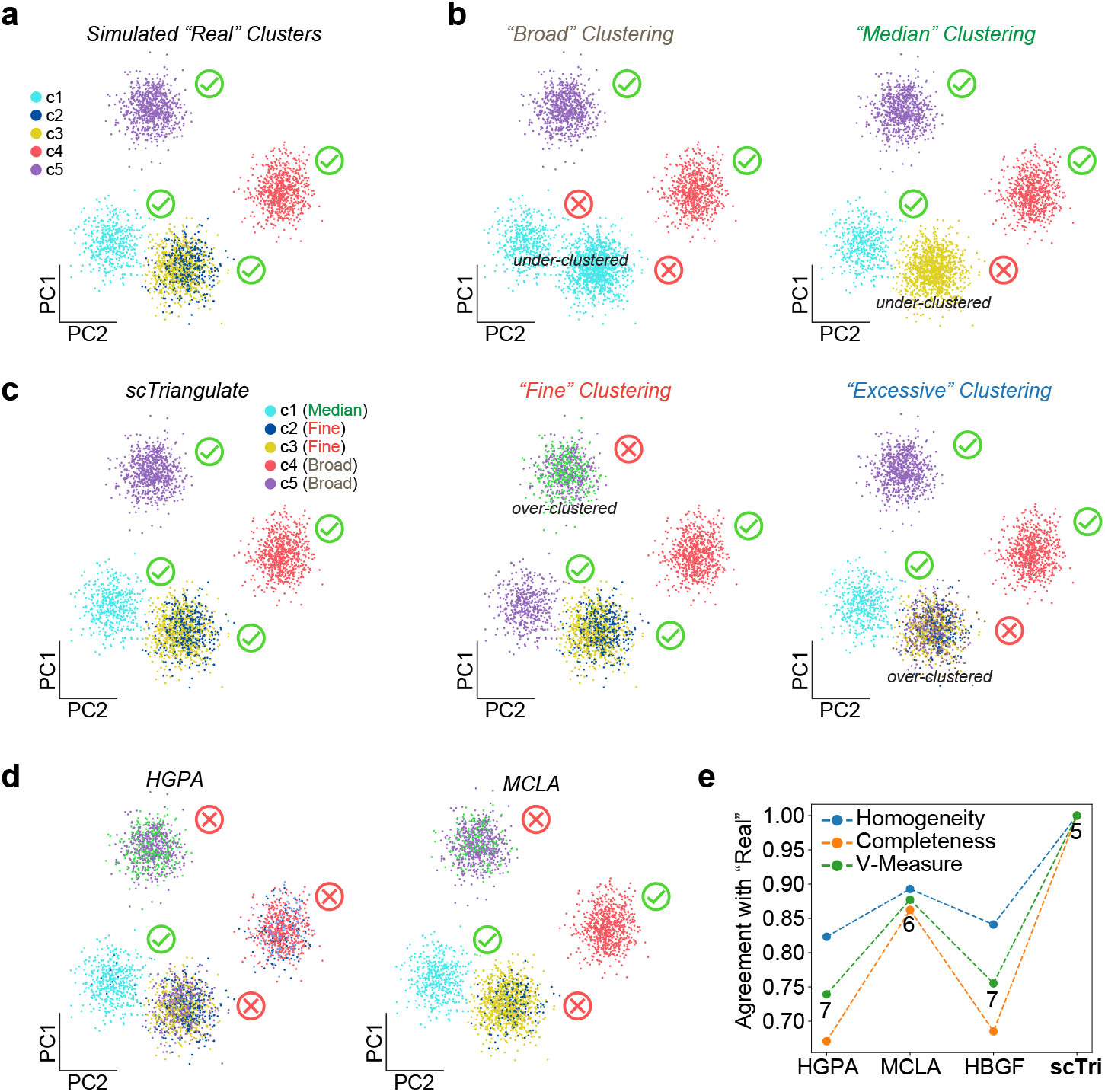
Heterogeneous scRNA-Seq clusters are uniquely defined by scTriangulate. a) As a ground-state truth to assess the specificity of predictions by scTriangulate, we simulated scRNA-Seq data for three highly distinct simulated populations (c1, c4, c5) and two subtly different subclusters (c2, c3) (Methods). b) To predict such clusters, we produce four cluster annotations for the same cells in panel a, with different granular cluster definitions (broad, median, fine and excessive). These include the noted under-clustered and over-clustered populations, which are not supported by gene expression differences. No individual resolution intentionally mirrors ground-state truth. Green checkmark = matching cluster to simulated. Red X = non-matching cluster to simulated. c) scTriangulate results using default parameters. The “resolutions” in which each cluster was selected from, is indicated in the cluster legend. d) Example ensemble clustering outputs from HyperGraph Partitioning Algorithm (HGPA), Meta-Clustering Algorithm (MCLA) and Hybrid Bipartite Graph Formulation (HBGF) applied to the four input resolutions. e) Benchmarking of the cluster assignments using three performance metrics (Homogeneity, Completeness, V-Measure) for three ensemble clustering techniques and scTriangulate. The number of clusters is indicated below V-Measure.

### scTriangulate finds novel common cell-states across uni- and multimodal platforms

To determine whether scTriangulate multimodal integrated results: 1) correspond to well-described cell states, 2) have improved accuracy over alternative approaches, and 3) reveal new discrete cell populations, we first applied it to several independent human immune single-cell datasets assayed with four distinct approaches: scRNA-Seq (RNA), CITE-Seq (ADT+RNA), multiome (ATAC+RNA) and TEA-Seq (ADT+ATAC+RNA). For the analysis of snATAC-Seq, scTriangulate adopts a modified version of epiScanpy ^17^ to collect peak-level information for the ATAC cell clusters. For ADTs, scTriangulate uses Centered Log Ratio (CLR) normalization ^1^. For these analyses, we consider a spectrum of software resolutions for each modality in the game. For TEA-Seq, our initial clustering considered 9 annotation sets from all three modalities and three Leiden resolutions, resulting in 11-38 clusters per resolution that span 203 possibilities (**Supplementary Figure 3a**). In contrast, projection of labels from a PBMC reference atlas (Azimuth) assigned labels for 14 cell populations, with greater than 10 cells (**Figure 3a**). Using scTriangulate, we obtain 19 final cell populations composed of source clusters from all considered modalities and resolutions (**Figure 3b,c**). Considering the Shapley associated confidence of the final clusters finds that nearly all cell populations are high confidence, with novel scTriangulated proposed populations (not in Azimuth) resulting in lower confidence but stable predictions (**Figure 3d**). Importantly, scTriangulate finds all predicted immune cell-types assigned by a comprehensive multimodal reference PBMC donor atlas ^5^, including challenging to detect and rare immune populations in blood (NK CD56+ bright, B-memory and HSCP). Notably, scTriangulate finds that distinct modalities (gene accessibility, transcriptomic or cell-surface features) often distinguish highly related cell populations. For example, B-memory cells were resolved primarily by ATAC-Seq features, CD8 naive by ADT and B-naive by RNA (**Figure 3e**). In addition to these well-defined populations, scTriangulate identifies new clusters (CD4 and B-cell), frequently defined by gene accessibility or cell-surface proteins. For example, we observe subdivision of CD4+ T-cells by the unique accessibility of a critical early T cell progenitor regulatory factor (*ZBTB17* ^18^), which was not resolved at the level of gene-expression (**Supplementary Figure 3b**). These and other novel subdivisions, were further supported by cell surface protein expression in T-cells (CD45RA, CD95, KLRG1) (**Figure 3f**). To benchmark these predictions, we compared cluster assignments for individual modality clustering solutions, multimodal integration and ensemble clustering for a range of software resolutions to the Azimuth reference (**Figure 3g**). Here, scTriangulate was found to have improved performance over all evaluated solutions and resolutions, including state-of-the-art multimodal integration (Seurat-WNN, totalVI) and ensemble approaches (MCLA, HGPA, HBGF).

**Figure 3.**
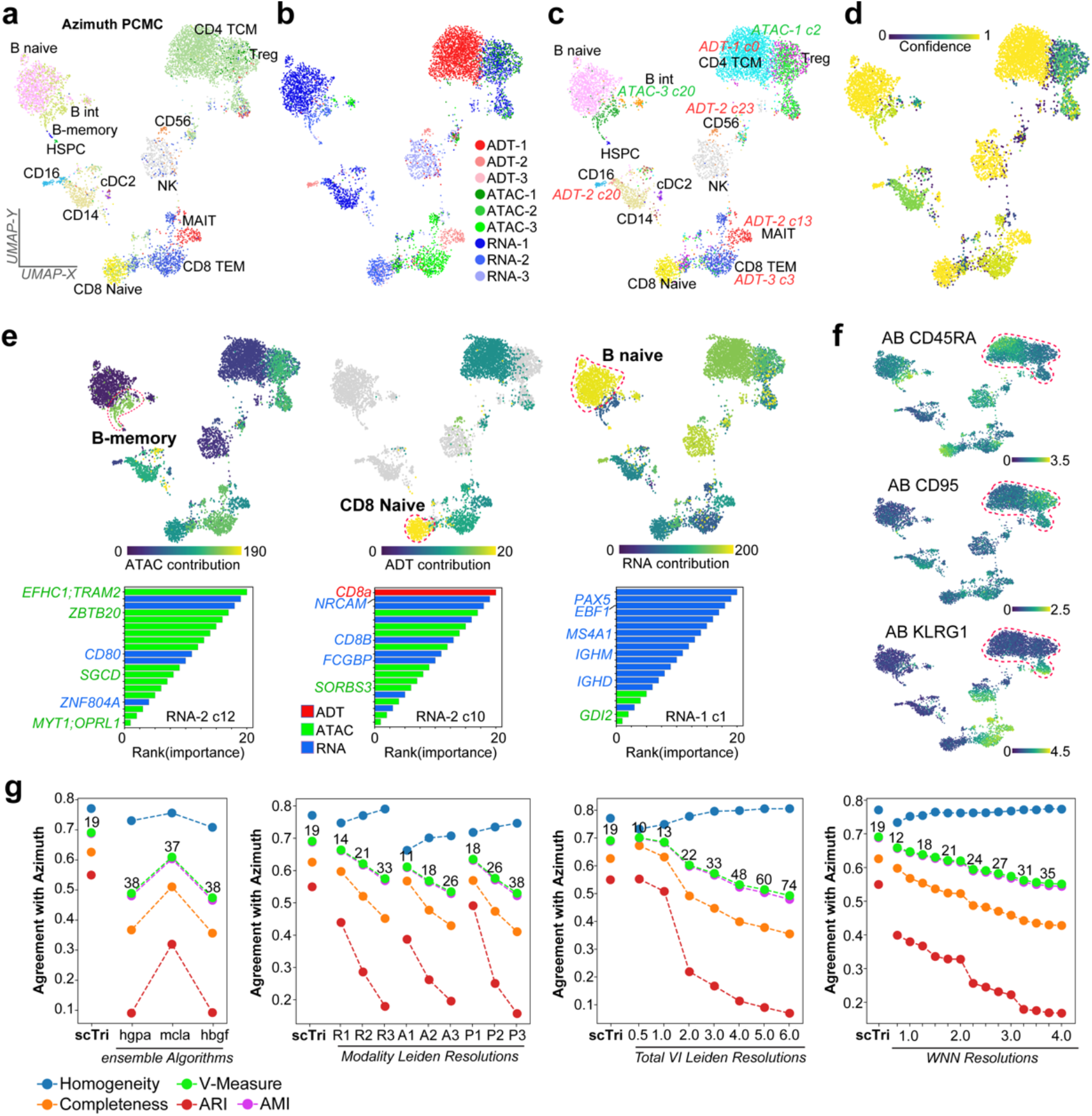
scTriangulate identifies discrete modality specific populations with improved performance over alternative methods. a) Supervised assignments from prior CITE-Seq cluster PBMC predictions (Azimuth). b) Integration of nine independently obtained clustering predictions from tri-modal GEX+ADT+ATAC (TEA-Seq) in donor PBMCs. Each color indicates a distinct modality-specific clustering resolution (1-3), with the winning scTriangulate clusters shown (UMAP based on ADTs). c) Clusters specifically derived from ADT ATAC cluster resolutions are labeled in the plot. d) Confidence of each scTriangulate-defined cluster. e) The selective contribution of each indicated modality is overlaid on the UMAP (top panels), based on the frequency of associated features among the top-20 markers of each final cluster (scTriangulate visualization). In the bottom panel, the top-ranked features for example clusters are indicated by their modality (color), with example features denoted. f) Novel T-cell subsets evidenced by cell-surface ADTs. g) Agreement with Azimuth PBMC reference label assignments measured by Homogeneity, Completeness, and V-Measure, for each individual or integrated set of clustering solutions (single-modality Leiden, multimodal Seurat WNN or TotalVI, ensemble clustering). Ensemble clustering was performed using 9 independent Leiden clustering results derived from the three independent TEA-Seq modalities (RNA, ATAC, ADT), using three popular consensus algorithms; HyperGraph Partitioning Algorithm (HGPA), Meta-Clustering Algorithm (MCLA) and Hybrid Bipartite Graph Formulation (HBGF). The number of clusters produced by each indicated resolution are displayed above the V-Measure.

To assess the reproducibility of scTriangulate on independent datasets, we generated CITE-Seq on peripheral mobilized human total nuclear cells (TNC) from two donors (donor 1 and 2). We again consider a set of clustering solutions varied across resolutions and modalities (**Supplementary Figures 4a**). scTriangulate successfully detected nearly all reference-defined cell populations in both samples, except for rare CD56+ high NK cells and plasmablasts in the first donor and intermediate B-cells in the second donor (**Supplementary Figures 4b-g**). In the case of CD56+ high NK, this population was detected in the triangulation of donor 1 and 2 (ADT-defined), but was “pruned” for donor 1, as only a fraction of the original cells was retained in the final result using default thresholds. Beyond these reference Azimuth populations, scTriangulate uncovered distinct subtypes of monocytes and MAIT cells, defined by the unique combination of RNA and ADT features (**Supplementary Figures 4h-j**). Importantly, these distinct subsets of MAIT cells, previously proposed to have increased responsiveness to innate cytokine stimulation, could only be derived through the integration of ADT and RNA clusters, as no source resolutions identified these populations ^19^. Similar to TEA-Seq, scTriangulate of both donors again had improved performance against all individual modality resolutions (**Supplementary Figures 4k**).

While these scTriangulate results show strong concordance with previous annotations, when applied to multiome RNA+ATAC PBMC this approach yields many high-confidence novel cell populations directly attributed to ATAC-seq features (monocytic, T cell) (**Figure 4a-c, Supplementary Figure 5a-c**). As such, relative performance of scTriangulate by V-measure and Completeness, was lower than a subset of independent clustering predictions (**Supplementary Figure 5d**). However, comparison of the molecular markers in these novel clusters indicates they represent: 1) classical, intermediate, and inflammatory CD14 monocytes^20^ (**Supplementary Figure 5e**), 2) effector memory, naïve and central memory CD4 T-cell subsets^21,22^ (**Figure 4d,e**) and 3) *CASC15*-defined innate lymphoid cells (ILC) ^23^ (**Figure 4e**). In addition to being evidenced by distinct and common ATAC and RNA features, these same novel monocytic populations and associated markers were evidenced in the CITE-Seq (**Supplementary Figure 5f**), demonstrating the scTriangulate predictions are highly sensitive and specific across diverse software resolutions and modalities.

**Figure 4.**
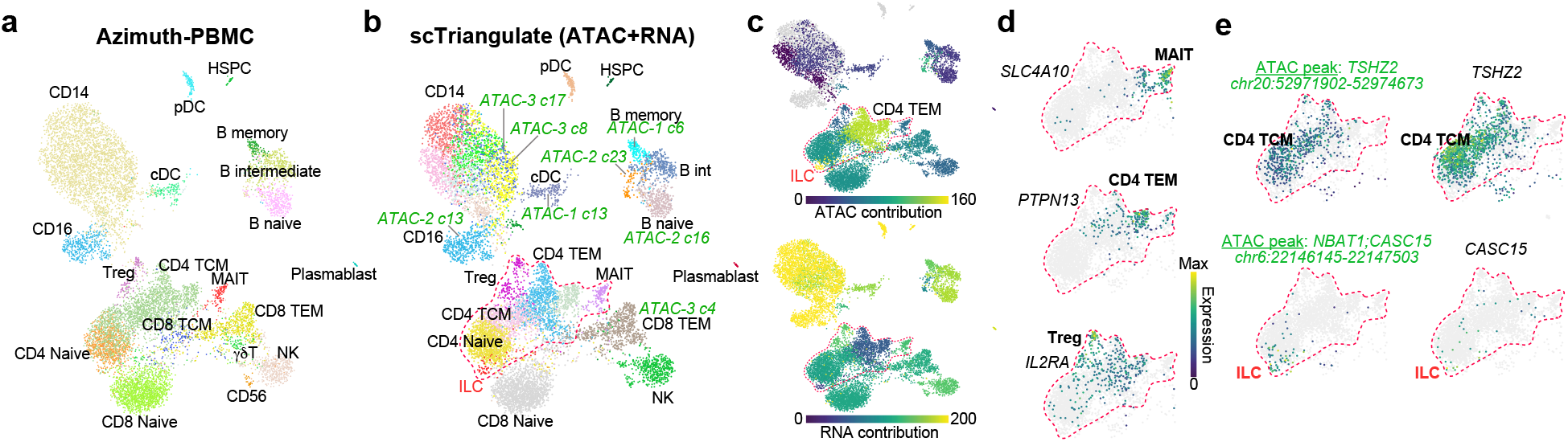
Multimodal integration identifies rare and novel cell-states distinguished by DNA-accessibility. a) UMAP of multiome PBMC (RNA+ATAC) following Azimuth assignment of cell population identities. b) scTriangulate final clusters after integration of multiple RNA and ATAC clustering resolutions. c) The selective contribution of each indicated modality is overlaid on the UMAP, based on the frequency of associated features among the top-20 markers of each final cluster. d,e) Visualization of the top-multiome RNA or ATAC markers for indicated rare or novel T-cell subsets. e) Correspondence of the abundance of unique ATAC-Seq markers (e.g., *TSHZ2* marks canonical naïve/CM ^24^) with RNA expression of the same localized gene.

To determine whether unimodal scRNA-Seq alone could identify similar populations defined by multimodal analysis, we only considered scRNA-Seq for different software settings (resolutions) (**Supplementary Figure 6a**). As expected, the optimal solution defined by scTriangulate does not specifically correspond to a single individual Leiden resolution, but rather a mixture of multiple resolutions (**Supplementary Figure 6b,c**). Further, scTriangulate confidently identified 22 out of the 25 Azimuth cell populations with >10 aligned cells (**Supplementary Figure 6d,e**). Notably, these “missed” populations (B intermediate, CD4 Proliferating, NK CD56 bright) were not identified from the finest tested resolution and thus could not be considered in the game. scTriangulate predictions included subtle population differences in diverse T-cell subsets, such as the correct split between Azimuth predicted CD8

T-effector memory (TEM) and CD8 T-central memory (TCM) cells. Such predictions result from improved stability scores and Shapley value for each all evaluated metrics (Reassign, SCCAF, TF-IDF) (**Supplementary Figure 6f,g**). Beyond these Azimuth predictions, which only finds a single CD14 monocyte population, scTriangulate finds the same novel subdivisions of classical, intermediate, and inflammatory CD14 monocytes, as scTriangulate’s CITE-Seq and multiome analyses (**Supplementary Figure 6h**). Finally, to ensure the default parameters applied in this study are robust for diverse multimodal study designs, we compared them to alternative parameters, including alternative ranking strategies, cluster pruning options and decision strategies for annotation importance for all evaluated blood datasets. These evaluations demonstrate that scTriangulate defaults result in optimal multimodal clustering integration (**Supplementary Figure 7**). Hence, scTriangulate’s predictions are highly accurate relative to prior references while revealing reproducible novel populations.

### Automated cell atlas aggregation across studies

Motivated by the challenge of multimodal integration, we asked whether scTriangulate can systematically integrate and even replace cell atlas expert curation. Current protocols for cell-atlas construction require complex dataset integrations, multiple rounds of re-clustering with different resolutions and tools, alignment to different curated references and extensive manual curation by domain experts. In the lung, diverse clustering algorithms, single-cell technologies, cluster resolutions and extensive manual curation have been used to identify and characterize dozens of distinct cell types. Indeed, projecting 142 labels from four lung atlas-level studies onto a single dataset ^25^ (**Methods**) illustrates significant diversity and conflicts in cell-type heterogeneity, cluster boundaries and labels ^26–28^ (**Supplementary Figure 8a**). To resolve these conflicts, we applied scTriangulate, which produced a single high-confidence and comprehensive lung atlas, borrowing the best cell-type definitions from independent studies (**Figure 5a-d**). For example, scTriangulate was able to subdivide dendritic cells (DCs) and alveolar macrophages into more informative sub-populations than present in any single-reference lung atlas (**Figure 5e**) ^29^. Comparison of scTriangulate predictions to a cross-tissue mononuclear phagocytes atlas called MNP-VERSE, supports our finding of expanded DC and macrophage heterogeneity in the lung (**Supplementary Figure 8b**) ^30^. While in many cases, more granular cluster predictions are rejected by scTriangulate (e.g., ncMono and cMono in Adams) (**Supplementary Figure 8c**), we cannot rule out the possibility that such populations will be better resolved in larger patient cohorts or considering additional modalities. Further, as the naming of populations will often vary between studies (example in **Supplementary Figure 8d**), expert curation of the scTriangulate integrated labels will always be required. To further assess the quality of scTriangulate cluster predictions and names, we compared these to two independent integrated lung atlas annotation efforts (Human Lung Cell Atlas (HLCA) and CellRef, **Methods**) (**Figure 5f** and **Supplementary Figure 8e**). In addition to identifying confirmatory support for many of scTriangulate’s subdivisions (epithelial, endothelial, myeloid, B-cells) and names, scTriangulate had improved agreement with independent lung atlases over those obtained with ensemble clustering (**Figure 5g** and **Supplementary Figure 8f**).

**Figure 5.**
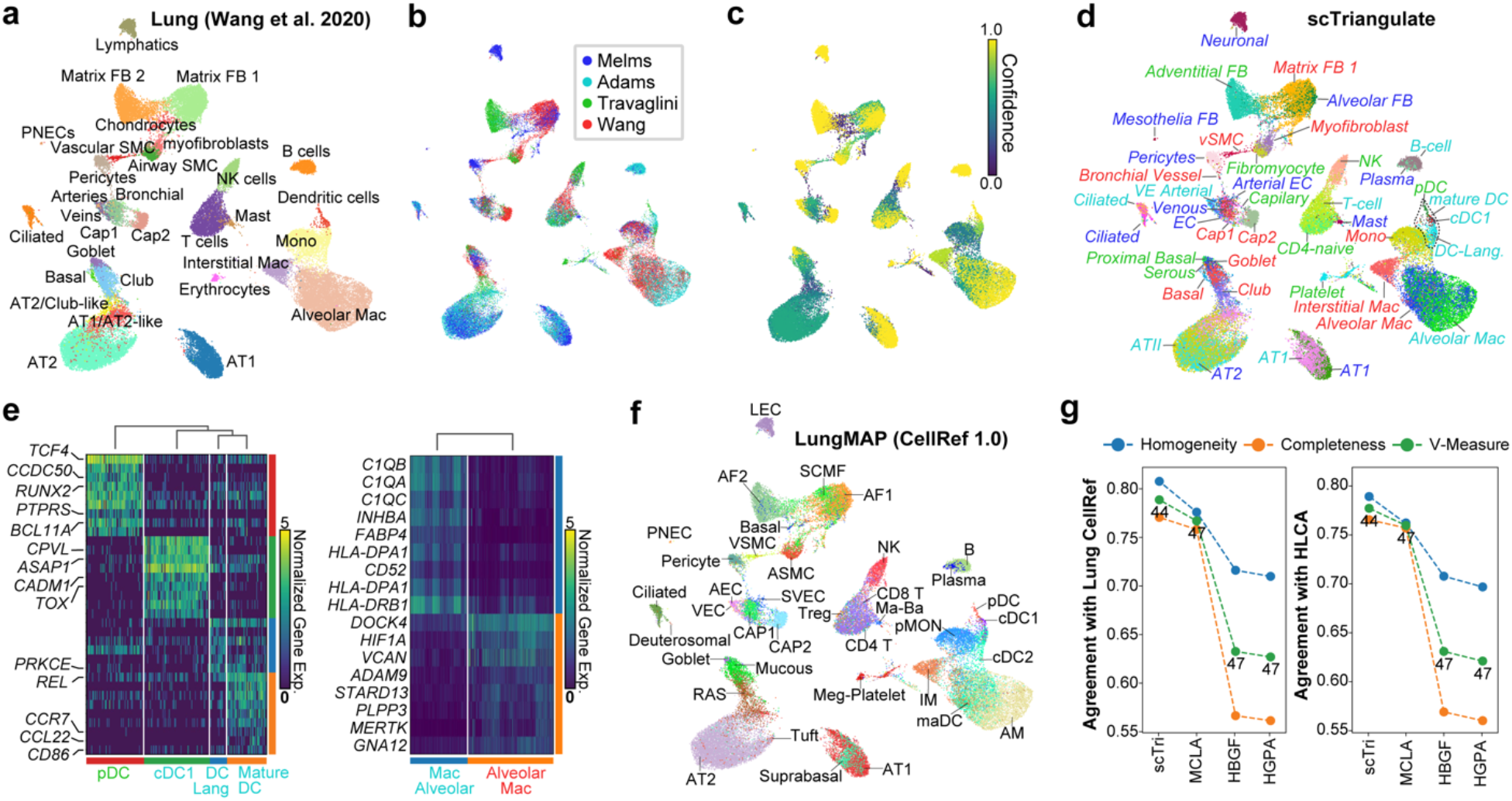
Automated game-based integration of cell-atlases. scTriangulate of 4 competing lung cell atlases projected onto snRNA-Seq from 9 donors (Wang et al. 2020). a) Wang et al. source annotations and scRNA-Seq. b) scTriangulate winning clusters after integrating labels from three other lung atlas studies projected onto Wang (Melms, Adams, Travaglini) (**Methods**). c) The confidence of each scTriangulate-defined cluster based on the “winning-fraction” of cells in each cluster (**Methods**). d) scTriangulate final label assignments, corresponding to panel b. e) Heatmap displaying the top discriminating dendritic (left) and macrophage (right) cell markers for Wang et al. cluster subpopulations determined from scTriangulate (literature implicated gene symbols displayed). The cell (d) or label (e) color corresponds to the source dataset annotations in panel b. f) Cell annotations from human LungMAP CellRef labels (Azimuth projected - finest level) onto the Wang snRNA-Seq data. g) Agreement with Azimuth CellRef (left) or HLCA (right) reference label assignments measured by Homogeneity, Completeness, and V-Measure for scTriangulate and ensemble clustering.

To determine whether scTriangulate can resolve more granular cell populations in an established cell atlas, we integrated diverse clustering annotations (ICGS2, Seruat3, Monocle3, curated) in an existing human bone marrow cell atlas composed of over 100,000 cells from eight donors^31^ (**Supplementary Figure 9a-d**). Importantly, the prior atlas proposed discrete hematopoietic stem and progenitor populations which have been previously described in a small subset of the original captures (<10% of cells). Importantly, these rare populations were invariably captured by these independent clustering solutions ^8^. While analysis with only a single TF-IDF score failed to resolve certain transitional states (e.g., HSC-cycle, MEP, LMPP), these were readily identified using the default options, which distinguish more granular cell-states (**Supplementary Figure 9d,e**). Evaluation of the default scTriangulate clusters with individual source annotations resulted in a greater agreement with the original author annotations (**Supplementary Figure 9f**). In addition to transitional cell states, scTriangulate was able to identify prior literature curated dendritic, monocytic and erythroid subsets, not defined by the original authors (**Supplementary Figure 9g-i**).

### Resolving clonal impacts from genotype aware single-cell RNA-Seq

A new and increasingly important form of multimodal single-cell genomics is the analysis of clonal genomic architecture and transcriptomic impacts. While the clonal architecture of cells from a patient can be identified with a growing number of multimodal protocols, no current integrative methods exist to determine how mutations contribute to neoplastic states. Therefore, we applied scTriangulate to single cell gene expression with linked gene-mutation calls from a myeloproliferative neoplasm (MPN) patient ^2^. Here, a small number of mutations were selectively detected from the same 10x Genomics scRNA-Seq MPN library, through sub-library amplification using the Genotyping of transcriptomes (GoT) protocol. Using GoT, the original authors identified wild-type hematopoietic cells and three neoplastic clones (*SF3B1, SF3B1*+*CALR*, or *SF3B1*+*CALR*+*NFE2*) (**Figure 6a,b**). First, scTriangulate integration re-assigned cells (*SF3B1* single mutants) to potentially more accurate clonal populations, as reflected by their more similar transcriptomes (**Figure 6d**). Such reassignments are crucial, due to incomplete genotyping which is inherent in methods such as GoT. Next, scTriangulate revealed *SF3B1*+*CALR*+*NFE2-*clone-induced stable cell states within most cell types sampled (e.g. HSC, MultiLin, etc), but the *SF3B1* and *SF3B1*+*CALR* clones could only be distinguished in the MEP and early-ERP trajectory (**Figure 6c**). This suggests that the genomic impact of *CALR* mutation within MEP and early-ERP is unique, or that CALR impacts in earlier hematopoietic cells is eclipsed by the impact of *SF3B1*.

**Figure 6.**
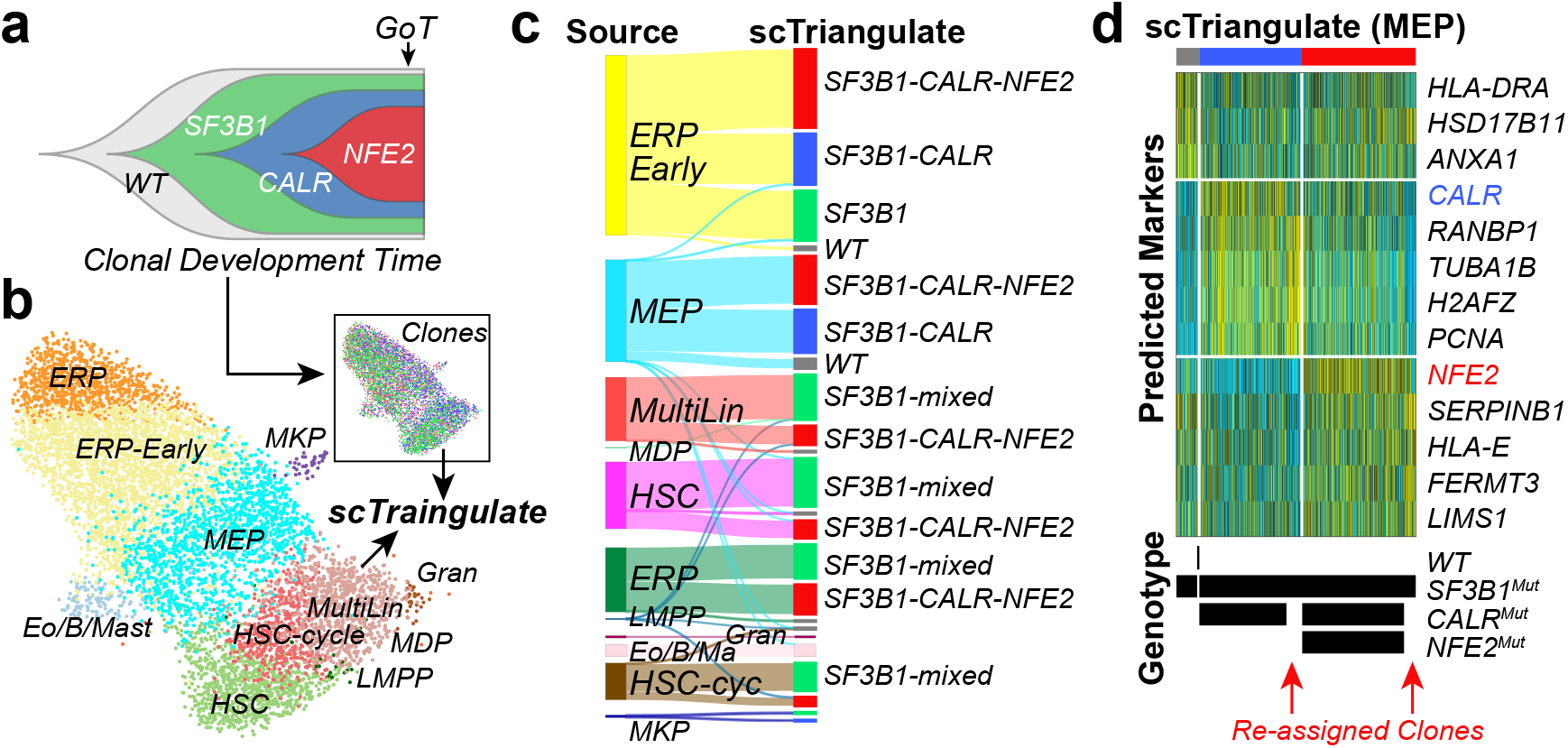
Detection of cell-state-specific clonal impacts from Genotyping-Of-Transcriptomes. a) Fish-plot displaying the clonal evolution of a patient with one (SF3B1), two (SF3B1-CALR) or three (SF3B1-CALR-NFE2) causal MPN mutations. GoT (arrow) indicates when single-cell profiling and genotyping were performed. b) MPN single-cell clusters assigned by supervised assignment (cellHarmony) relative to healthy bone marrow CD34+ progenitors, with clonal architecture of the cells indicated in the inset. c) Sankey diagram of scTriangulate transcriptionally coherent cell populations, considering gene expression and clonal genotype. d) Expression heatmap of final scTriangulate clusters for MEP, indicating the original predicted genotypes from GoT and scTriangulate reassignments.

### Alternative splicing distinguishes subpopulations of leukemic blasts

Similar to clonal variation in cancer, alternative splicing is an established critical driver of oncogenesis in the presence and absence of spliceosome mutations ^32^. To determine whether scTriangulate, in conjunction with sparse splice-sensitive droplet scRNA-Seq, can unambiguously identify splicing-defined cellular subtypes, we profiled pediatric leukemia progenitors using the 10x Genomics 5’ platform. Since the definition of novel cell splicing-defined cell populations has not been previously successful due to the extremely sparse exon-exon junction data, we developed a new strategy to identify candidate splicing-defined subclusters using gene expression-defined hematopoietic progenitor clusters (**Figure 7a, Methods**). scTriangulate integration of gene expression- and splicing-defined populations identified new leukemia blood progenitor populations that subdivide known cell-types (**Figure 7b-e, Supplementary Figure 10a-d**). For most cell-types, only one splicing-defined subcluster was retained, with the others effectively merged, based on their stability. These splicing perturbations alter the composition of numerous hematopoietic stem and progenitor (*KLF2, GATA2, GATA1, SPINK2*) or leukemia (*LAT2* and *NUCB2*) regulatory factors. Observed events were readily confirmed by aggregate SashimiPlot visualization (**Figure 7f** and **Supplementary Figure 10e**). Importantly, such clusters could not be resolved through multimodal integration of the underlying splicing and gene expression data using Seurat WNN (**Supplementary Figure 10f,g**).

**Figure 7.**
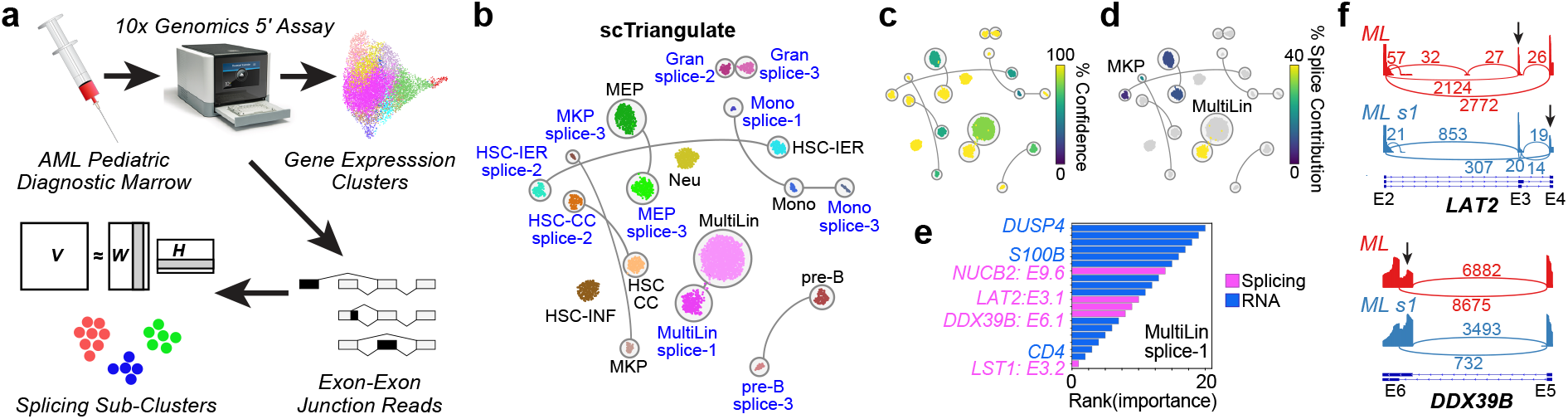
Resolution of novel splicing-defined cancer cell-states. a) Overview of the strategy for identifying initial splicing-defined subsets, from a diagnostic pediatric AML bone marrow sample profiled using the 10x Genomics 5’ library chemistry. Initial clusters were defined using the software ICGS2 from gene expression measurements and k=3 sub-clusters from non-negative matrix factorization (NMF) analysis of percent-spliced-in (PSI) values. b) UMAP of AML cells from neighborhood component analysis of AML gene expression and splicing, guided by scTriangulate analysis final “pruned” labels. Remaining stable splicing-defined sub-clusters (NMF clusters 1, 2 or 3) are indicated in blue. c) The Shapley-associated confidence of each scTriangulate-defined cluster. d) The selective contribution of splicing is overlaid on the UMAP. e) Top-ranked markers (splicing and gene) in the MultiLin (ML) splicing-defined sub-cluster (s1). f) Selected splicing event Sashimi-Plots comparing ML denoted scTriangulate sub-clusters.

## DISCUSSION

A major goal of single-cell genomics is the accurate isolation and functional characterization of predicted cell populations. Prior to validation, it is essential to have statistical confidence in the underlying predictions, with consideration of diverse possible alternatives. Given hundreds of existing approaches, rather than argue for one solution as inherently better, we argue that optimal clustering decisions require the integration of diverse cellular annotations from independent studies (i.e., reference-classification), multimodal measurements, distinct clustering algorithms and software settings to identify the most confident, as opposed to the most common (ensemble) predictions. scTriangulate represents a fundamentally distinct approach for integration, that is fast, accurate and which can be effectively scaled to any number of new modalities or stability metrics (**Supplementary Figure 11**).

As demonstrated in our evaluation studies, prior known and novel populations defined by scTriangulate are frequently informed by distinct-modalities, can be consistently identified between different multimodal single-cell technologies and have improved correspondence to highly curated references than prior multimodal integration approaches. Further, our results show that the integration of prior obtained annotations from lung or bone marrow identifies improved cellular identities that can only be obtained from the combination of multiple independent sources. As such, scTriangulate has the strong potential to automate cell atlas creation through the integration of different proposed annotations and new clustering results. These can include new populations which arise from partially overlapping annotations from different sources. An important remaining challenge of such analyses is the naming of populations, which remains largely non-standardized (see **Supplementary Figure 8**).

In the context of clonal analyses in cancer, we show for the first-time multimodal integration of gene expression and alternative splicing to illustrate splicing-defined clonal subsets in cancer as well as mutationally-defined clonal subsets that result in dominant transcriptomic impacts. While our current study was limited to the measurement of select somatic mutations within a tumor, our approach can logically be extended to the unbiased analysis of hundreds or thousands of genomic variants, identified at the DNA or RNA-level using newly emerging single-cell multimodal genomics protocols^33^. Such analyses are likely to be crucial to differentiate cancer clones *in vivo* based on their gene, long-read, splicing, cell surface or epigenomic readouts. Hence, as the number of single-cell modalities increase, it will be necessary to not only integrate cellular heterogeneity from each molecular readout, but to combine such predictions with reference cell annotations from existing studies that involve diverse computational tools.

While the analytical settings applied in these studies were large programmatic defaults, it is likely these settings will not be optimal for all study designs. For example, the identification of rare bone marrow progenitor populations required the use of multiple TF-IDF scores to weight the selection of rare transitional cell-states with fewer unique expressed marker genes. Further, the integration of cluster predictions from dozens of sources (e.g., resolutions, modalities, reference annotations), will significantly increase runtime. For these reasons, the software includes intelligent defaults, alternative options for stability metric inclusion and flexible parameters (e.g., pruning threshold), that allow users to customize the outputs of scTriangulate out of the box. For example, the program will automatically substitute Shapley selection of final clusters with combined rank-based selection, when over 15 annotation sets are provided to reduce runtime. scTriangulate can be further customized beyond such options, to include new stability metrics or alternative rank-integration approaches.

Looking forward, we anticipate extension of this approach beyond single-cell genomics to more complex settings, such as spatial transcriptomics, in which the notion of molecular modalities can be extended to spatial position within a tissue, cellular niches, subcellular localization, cellular interactions, cell morphology, and other qualitative and quantitative readouts enabled through image-based analyses. scTriangulate is extendable to such approaches and already possesses an interface to the python spatial transcriptomics analysis package Squidpy ^34^. In addition to the analysis of new technologies, it will be important for this and similar approaches to support the integration of multiple samples of the same modality and even across different single-cell multimodal or spatial technologies. We aim to support such features in the future, through the incorporation of innovative new computational strategies.

## ACKNOWLEDGMENTS

We would like to thank Kang Jin, Balaji Iyer and Daniel Schnell for their valuable feedback on the approach and manuscript. We also thank Kelly Rangel in the CCHMC Gene Expression Core for generating CITE-Seq libraries and Andre Olsson for his assistance. This work was partially supported by the National Heart Lung and Blood Institute (R21HL150678 to H.L.G and N.S., U24HL148865), National Institute of Diabetes and Digestive Health (RC2DK122376 to H.L.G. and N.S.), the Chan Zuckerberg Pediatric Network for the Human Cell Atlas (N.S.) and the Cincinnati Children’s Pediatric Cell Atlas Center (N.S. and H.L.G).

## AUTHOR CONTRIBUTIONS

G.L. implemented the method and performed evaluations. B.S. generated the CITE-Seq data. N.S., G.L., H.L.G., H.S. and V.B.S.P designed the study and wrote the manuscript.

## CODE AND DATA AVAILABILITY

scTriangulate is available as a Python3 package (https://pypi.org/project/sctriangulate). The source code and docker container are available at (https://github.com/frankligy/scTriangulate). The scripts and data for reproducing the results are available at (https://github.com/frankligy/scTriangulate/tree/main/reproduce)

The peripheral blood CITE-Seq and AML scRNA-Seq processed results, scripts, relevant outputs and metadata has been deposited in Synapse (https://www.synapse.org/#!Synapse:syn26320566)

## METHODS

### scTriangulate Algorithm

scTriangulate is organized into 5 principal steps (**Figure 1**). In the first step, counts or normalized data matrices are loaded in the software for QC, filtering, and scaling (optional). This step may also include running Leiden clustering sequentially through scTriangulate when no prior annotations exist. In the second step, cluster stability is computed for all input annotation-sets. In the third step, a Shapley Value is computed to measure the importance of each annotation set and assign each cell to a supplied or scTriangulate computed set of labels. In the fourth step, the final annotation assignments are pruned according to the fraction of final versus original cell-cluster assignment (user-defined) and re-classified using a nearest-neighbor strategy. Finally, these results are reported to the end-user using a series of interactive or static (pdf) visualizations to explore locally or over the web. Additional details and tutorials for scTriangulate are provided online with the software.

#### Step 1: Annotation assignment and quality control

Input data matrices and options are supplied to the software to enable automated processing of a dataset without existing annotations (i.e., clusters) or with externally produced annotations. This is performed using the preprocessing module wrapper function scanpy_recipe. scTriangulate begins with two data matrices **X** and **O**, where **X** ∈ **R**^**I×F**^ denotes the expression matrix and **X**_**if**_ is the scaled expression value of feature **f** in cell **i**, with the total number of features as **F** and the total number of cells as **I**. The features denote a gene, ADT, ATAC-Seq peak, or custom modality-specific features. By default, scTriangulate assumes that the input expression data is log scaled values (counts per 10,000 reads per cell + 1). Let **O** ∈ **R**^**I×A**^, where **O**_**ia**_ denotes the cluster label of cell **i** in annotation **a**, with the total number of annotations as **A**.

#### Step2: Inference of cluster stability

To decide which cluster assignments are optimal, the software computes a series of metrics to gauge the reliability of each cluster for each annotation. scTriangulate uses four embedded scores: the **reassign** score, the **TFIDF10** score, the **TFIDF5** score and the **SCCAF** score. These default scores can be augmented by users with custom metrics to suit to either promote or disqualify clusters with certain properties (doublet, cell cycle, metabolic pathway, etc). Here we describe the four default scores used in the program:

##### Reassign Score

The reassign score quantifies the reliability of a cell-to-cluster assignment when compared to its own centroid (nearest neighbor distance). First, scTriangulate defines the top-ranked 30 markers using an integrative strategy considering both log-fold change (effect size) and a two-sided t-test p-value (significance) for each cluster versus all other cells. Here, the union of all discriminating features for all clusters are denoted by **L**. Next, a PCA analysis is performed on **L** to obtain a reduced dimensional representation **P**. Based on the discriminative features and the derived PCA space, scTriangulate reduces the dimension of the dataset from **F** to **P** and the centroid of each examined cluster is derived from these features, to produce a centroid matrix ∈ **R**^**P×C**^ where **C** is the total number of clusters in each analyzed annotation set. Specifically for ranking the markers, let **r**^**f**^_**pval**_**(c)** be the rank of feature **f** in cluster **c** based on the t-test p-value. The most significant feature will receive a rank=0. Then let **r**^**f**^_**lfc**_**(c)** be the rank of feature **f** in cluster **c** based on the log-fold change. Likewise, the features with the largest positive log fold change will receive rank=0. The combined rank will then be:

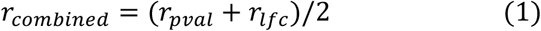

We define the function **NC(·)** to find the cluster label of the nearest centroid of each cell **i**, given the centroid matrix we defined above, and the Reassign score for each cluster **c** as:

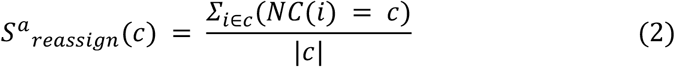

Where |**c**| denotes the size of cluster **c**, the reassign score effectively measures the fraction of cells in each cluster that can be reclassified to its own centroid. A higher value indicates greater stability.

##### TFIDF(n) Score

The TF-IDF score measures the number of exclusively expressed features in a cluster. This score is based on the observation that genes or other features that are uniquely expressed in one cluster typically demarcate the boundary of a potentially novel cell population. Unlike the reassign score which utilizes the expression values, the TF-IDF score focuses on expression frequency (zero and nonzero) as a binary value. To calculate the TFIDF score for a cluster, we first define the Term Frequency (**TF**) and Inverse Document Frequency (**IDF**) of each feature **f** in each cluster **c**. A pseudo value *∈* = 1e − 5 is added to avoid the numerical issues:

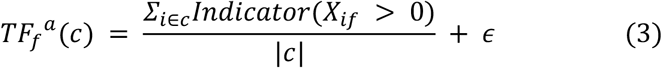

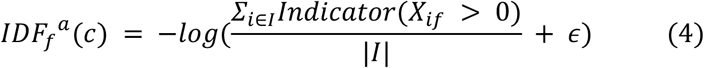

Where **I** denote cells in the whole dataset but **c** only contains cells in the selected cluster. Then we define the exclusivity, which is equal to the TF-IDF score of a feature in the cluster **c** as:

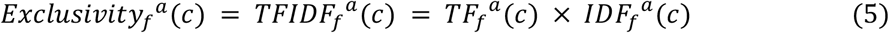

Next, this function ranks the features in each cluster based on their exclusivity in descending order. We can define the **TFIDF(n)** score of a cluster as the exclusivity value of the **n**th feature:

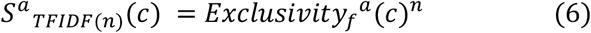

We consider **TFIDF5** and **TFIDF10** scores by default. As an example, *S*_*TFIDF*(5)_ (*c*1) means the TF-IDF score of the top 5th exclusively expressed feature in cluster c1. If a cluster expresses few unique marker genes (e.g., n=3) but is considered biologically valid, the user can add or replace the existing TF-IDF score with an alternative rank, such as TF-IDF1 to prioritize such features in the downstream Shapley analysis (**Supplementary Figure 2**).

##### SCCAF Score

We use the previously described Single Cell Clustering Assessment Framework (SCCAF)^11^ to serve as an added metric to gauge the biological reliability of the cluster. Different from the reassign score where we leverage the univariate markers from a two-sided t-test, SCCAF considers the marker definition challenge as a multivariate multi-class logistic regression problem, which inherently takes into account the interactions and dependencies between different features. While only the most discriminative features are considered for the reassign score, SCCAF considers all features. As it considers all features, SCCAF scores will not always be concordant with reassign and will differentially assess stability. In the original proposed model, the expression matrix and the Annotation will serve as the data **X** and the label matrix **O** respectively. After logistic regression model inference, a confusion matrix **Ψ** ∈ **R**^**C×C**^ will be produced. This matrix stores the number of times the model correctly and incorrectly predicts each instance’s cluster label. Using the confusion matrix, we define the SCCAF score of a cluster **c** as:

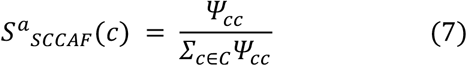

#### Step3: Shapley Value determination and cell-cluster assignment

The Shapley value is used in scTriangulate for fair credit allocation of the ranks derived from different stability metrics for each cell and annotation (“player”). Shapley applies coalitional iteration to compute the marginal contribution for each player. As this method tests the benefit of all possible combinations of players (cooperatively), in an iterative manner, compute time is relative to the number of players considered. Rather than directly computing annotation importance (Shapley) on the stability metrics, it is computed on the numerical sum of stability metric rankings using a process of coalitional iteration. Rank is determined considering all players within each possible coalition (all player permutations), hence, the rank of a stability metric is dependent on which players are considered in which coalition. Here, similar scores (within a tolerance threshold - aka offset) will be assigned the same rank, where ranks themselves are scaled in a “winner takes all” manner. The actual final Shapley value for a player is computed from the sum of the scaled stability associated ranks derived for each cell for a single player, considering all possible coalitions. This strategy has numerous conceptual and mathematical advantages over alternative strategies. For example, while simple rank-based annotation importance can be applied on the derived stability ranks (without coalitional iteration), fairly minor overall differences in the scores from one stability metric can override much larger differences in another and frequent ties between players. When different cluster annotations are considered equivalent (a tie), the software selects the smaller source annotation as the winner.

To compute, following the determination of stability metrics in **Step 2** for every single cell, a resulting matrix **D** ∈ **R**^**A×M**^ will be produced, where **D**_**am**_ denotes the calculated metrics of an annotation **a** under metrics **m**. For instance, if one selected cell has three conflicting annotations, namely label-1, label-2, and label-3. By default, matrix **D** will be of the shape 3 × 4 as scTriangulate compute 4 scores.

We consider each annotation here as one “player” in a cooperative game. To assess the relative importance of a player, the Shapley Value **Φ** has been proposed to guarantee fair credit allocation, which considers the amount of added contribution (marginal contribution) from each player (one label) to the collective performance of a team/coalition (multiple labels). With a proper Value function **V(T)** to represent the team’s performance, the Shapley Value of a label **a** can be computed as:

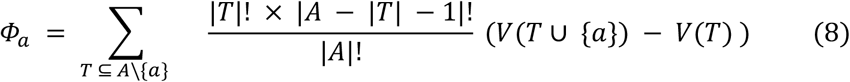

In brief, the Shapley Value iterates every possible formation of the teams and measures the marginal contribution of each player (label) to the team, normalized by the team size. scTriangulate defines the added performance (surplus) as below:

##### Algorithm 1

scTriangulate Computing (*V* (*T ∪ {a}*) *− V* (*T*))

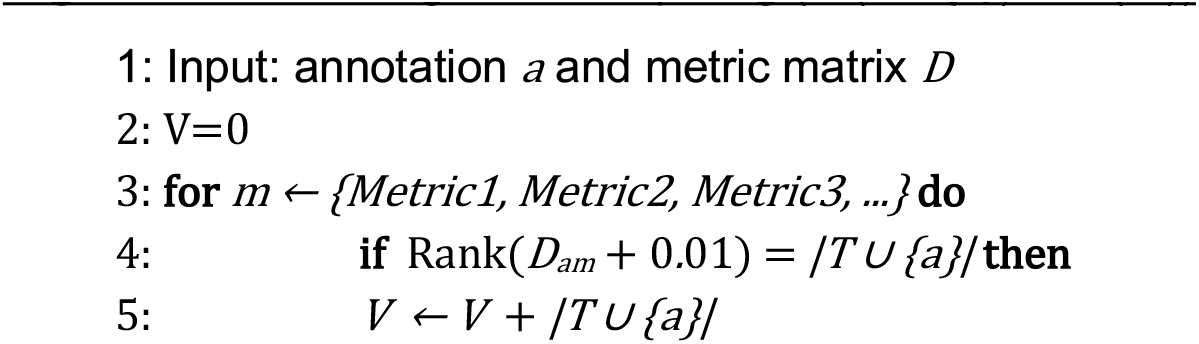

In the value function (**Algorithm 1**), 0.01 equals the default tolerance threshold or offset, to assign marginally close scores the same rank. The rationale for an offset is that two or more players will often be assigned a stability score for a given metric that are almost numerically equivalent but that result in different rankings. The use of an offset is inspired by General Linear Models (GLM), where an offset term is used to account for residual variation that is disproportionate and cannot be explained by the regression model itself. A fixed value offset is used as stability metric scores are in the range of 0-2 (adjustments may be required for new scores outside this range). We note that the optional use of an offset can improve overall performance in some but not all real-world benchmarking datasets (**Supplementary Figure 7b**). When computing the rank of a stability metric within a specific coalition, the final ranks undergo an optional adjustment. Specifically, the two existing adjustment options are “classic” and “all_or_none” (default). By default, the program re-scales all ranks for each cell and for each stability metric in a winner takes all manner (all_or_none), where only the top rank choice(s) retain its rank (all others set to 0). If multiple annotations have the top rank, then each of these annotations will be assigned the same non-zero rank. Hence, the all_or_none approach rank normalization approach only rewards a player if that cluster annotation has a higher score than all others for a given stability metric. The rationale for this option is that the best performers will rise to the top, producing more definitive ranking for downstream steps. Conversely, “classic” will retain all of the original ranks, post-offset consideration. The impact of these adjustment choices will impact the final results (**Supplementary Figure 7c**), most significantly, when simply ranking as opposed to Shapley is applied. After considering offset, the highest numerical value in **D**_**am**_ will receive the highest rank (scipy convention).

scTriangulate uses the **Φ**_**a(i)**_ to denote the relative importance of each label for each single cell **i**, and assigns this single cell **i** to the label with highest **Φ**:

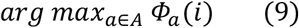

Since the computational complexity of Shapley increases exponentially as a result of exhaustive iteration of all coalitions, compute time becomes intractable with greater than 15 annotations. Hence, to reduce runtime, the program will default to an alternative solution purely based on rank, as a reasonable alternative to Shapley when the number of user-supplied conflicting annotations is greater than 15. Evaluation of pure ranking as the method of annotation importance demonstrates that it has generally equivalent importance (depending on the benchmarking dataset), when explicit ranking (classic mode) is used in place of all-or-none (winner takes all) rank adjustment (**Supplementary Figure 7c**).

To assess the overall stability and quality of Shapley selected annotations, the software includes the option to produce an average cluster Shapley value, analogous to the use of an Elo rating in chess. This quality metric is defined as the average of all cells’ Shapley value in all clusters, normalized by the number of players, and further normalized by the number of clusters. Since the Shapley calculation is an additive process, the more players considered, the higher Shapley value will be. This quality metric can be used to approximate which cluster is more stable even for clusters derived from different datasets.

#### Step 4: Results pruning and cell reclassification

As scTriangulate assigns stable populations at the level of individual cells, an optional pruning step was introduced to exclude cell populations with few cells retained relative to the source cell-to-cluster assignments. Here, we assume that if a large initial cluster of cells exists, but only a small minority of cells with that annotation are considered to be “stable”, that the overall predictions is more likely to be unreliable. Such stable minor subsets will often be due to technical artifacts (e.g., high mitochondrial expression, dying cells), but may represent a valid “activation” state of a parent cell type.

For this analysis we denote the new cell label based on the highest Shapley Value as **A**^**raw**^. We further filtered out unstable clusters based on two criteria:

1. winning fraction: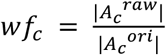 Stable clusters have a higher winning fraction of cells relative to the number of cells in the original cluster as compared to unstable clusters, which may win out purely by chance. By default, **wf**_**c**_ **< 0.25** will be labeled as unstable clusters. The winning fraction of raw clusters indicates the confidence that the program possesses for this prediction. In a benchmarking analysis of pruning thresholds ranging from 0-60%, we find that more pruning is predicted to always improve performance, but with minimal improvement beyond a threshold of 40% (**Supplementary Figure 7d**).
2. By default, |*A*_*C*_ ^*raw*^| < 10 will be labeled as unstable clusters. Specifically, scTriangulate only considers cell populations with a minimum number of cells (user-defined) for consideration of stability, since stability estimates in such populations are inherently less reliable due to potential outlier effects (abs_thresh parameter). Considering different possible thresholds for minimum cluster size (1-50 cells), the default threshold of 10 produced consistent reliable results for different evaluated datasets (**Supplementary Figure 7a**). All cells within the unstable clusters will be reassigned to their nearest neighbor cluster label.

##### Contributions of modality

The contribution of each modality {RNA, ADT, ATAC} in each cluster **c** is defined as a weighted sum of its markers (gene for RNA, antibody for ADT, peak for ATAC) in the top-20 marker list for the cluster **c**, the weight is the rank of the marker in the list such that the top1 marker will receive importance as 20. Specifically,

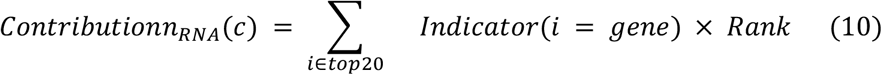

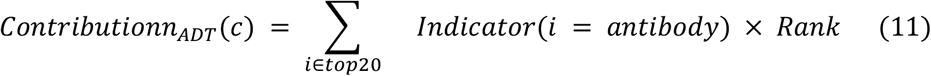

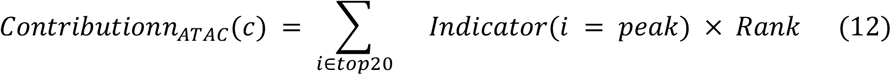

where **i** refers to each marker in the top-20 list, which can be either a gene, antibody or ATAC peak. Rank is the rank importance defined above.

#### Step 5: Results visualization

scTriangulate results can be exported en masse to an HTML archive to interactively explore marker expression, cell population boundaries, doublet cell predictions, quality control metrics, stability metrics, Shapley Values, modality contributions, and cell-population prediction confidence. These plots can be viewed as scatter plots (e.g., UMAP), heatmap, or bar charts. UMAP plots can optionally be exported from a supervised PCA analysis, considering the final scTriangulate clustering labels in the projection, using neighborhood component analysis (NCA). In addition, these plots can be individually exported from the software. Such visualization is an essential step to guide optimization of scTriangulate results to use alternative stability strategies (e.g., TFIDF5) or alter the cutoffs for pruning and reclassification in the stored scTriangulate object.

### scTriangulate Performance Evaluation

To assess scTriangulate’s ability to produce optimal uni-modal or multimodal clustering solutions, we compared its results to those to: 1) highly-used multimodal integration approaches, 2) ensemble clustering, and 3) individual clustering algorithms. As benchmarks we considered prior silver-standard curated atlas annotations (e.g., Azimuth) and synthetic scRNA-Seq data produced with Splatter. While the silver standard datasets are not definitive solutions, they represent significant annotation efforts where the original authors extensively analyzed different clustering solutions (algorithms, resolutions) and/or multimodal measurements (CITE-Seq).

#### Performance metrics

To assess relative agreement, we used three complementary cluster evaluation metrics, namely Homogeneity, Completeness and V-Measure. These three metrics were verified using Adjusted Mutual Information (ARI) and Adjusted Rand Index (ARI). Intuitively, Homogeneity denotes the extent to which each scTriangulate cluster only contains members of a single silver-standard reference cluster, or in other words, how homogeneous each scTriangulate cluster is. For annotations derived from low versus high clustering resolutions, it is expected that higher resolutions will increase homogeneity. Homogeneity is formulated as:

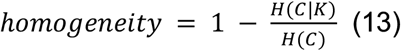

Where H(C) is the entropy of the silver-standard reference cell type C, and H(C|K) is the conditional entropy of the reference cell type C given the scTriangulate (or alternative clustering solution) cluster assignment K. Below, n is the total number of cells in the tested single-cell dataset, n_c_ is the number of cells for each reference class c, n_c,k_ is the number of cells that reference class c gets assigned to scTriangulate cluster k. Here, the mathematical notation is consistent with conventions applied in the sklearn python library.

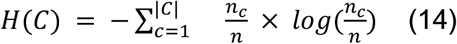

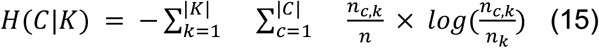

Conversely, Completeness aims to capture the extent to which all cells from each individual reference class are assigned to one single scTriangulate cluster. As expected, broader clusters derived from low resolution clustering will result in improved Completeness, as the chance of fully encapsulating a defined cell type is higher, unlike Homogeneity. Completeness is defined by:

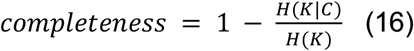

Where the H(K|C) denotes the conditional entropy of scTriangulate cluster assignment K given the reference cell type class C, n_k_ denotes the number of cells contained in each scTriangulate cluster k, n_k,c_ denotes the number of cells that scTriangulate cluster k comes from the reference cell type class c.

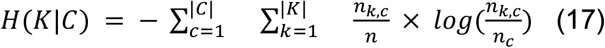

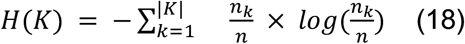

Clustering solutions in agreement with the reference should maximize Homogeneity Completeness score. V-measure is defined as the harmonic mean of both of these scores, which is formulated by Rosenberg and Hirschberg ^35^:

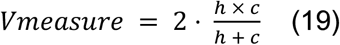

These metrics overcome potential pitfalls of alternatives, including no assumption of cluster structure and increased interpretability to diagnose whether a clustering resolution is too broad or too granular, which makes it suitable for evaluating single-cell clustering results, as previously demonstrated ^36,37^. While informative, as silver-standards are likely not comprehensive and often do not consider stability, results discordant with these assessments require stability, functional and/or literature verification to assess significance.

#### Simulation datasets

scRNA-Seq simulation datasets were generated using Splatter version 1.16.1 with *thesplatSimulateGroup* function. To assess the ability of scTriangulate to resolve different stable populations from different source annotations, a dataset was simulated with 3000 cells (*batchCells* = 3000). This ground-state truth dataset had 5 clusters (c1, c2, c3, c4, c5), where their size was determined by the parameter *group*.*prob* (0.23,0.15,0.15,0.23,0.24), in which each numerical value corresponds to the size of each simulated cluster. The extent to which two clusters are different is determined by parameter *de*.*prob* (0.15,0.15,0.15,0.2,0.4), where each numerical value corresponds to the fraction of genes that are differentially expressed in each cluster. We constructed four different annotations, which only partially overlap with the ground state solution. Specifically, we produced a “Broad” annotation that merges the above c1+c2+c3 (under-clustered), a “Median” annotation which merges c2+c3 (under-clustered), a “Fine” annotation that arbitrarily sub-divides c5 (over-clusters), and an “Excessive” annotation which arbitrarily sub-divides both c2 and c3. We apply the scTriangulate *lazy_run* function in version 0.9.0 on these four conflicting annotation solutions to derive the final cluster assignment (default settings).

To assess the impact of using one versus two TF-IDF scores, *batchCells* = 3000 was generated for 6 ground-state truth clusters (c1, c2, c3, c4, c5, c6) with their size determined by parameter group.prob (0.2,0.2,0.15,0.15,0.15,0.15). The extent to which two cluster groups are different is determined by *de*.*prob* (0.2,0.2,0.005,0.005,0.005,0.005), here we intentionally set the difference between c3-c6 to be subtle in order to assess the usage of multiple TF-IDF stability scores. Next, we constructed three different annotations, in which the “Broad’’ merged c3+c4+c5+c6 (under-clustered), the “Median” annotation combined c3+c4 and c5+c6 (under-clustered), and a “Fine” annotation which successfully identify the 6 simulated populations.

Here, we applied scTriangulate *lazy_run* function in version 0.9.0 using one TF-IDF (*add_metrics={}*) and two TF-IDF scores (*add_metrics={‘tfidf5’}*) on these three conflicting annotations to derive the final cluster assignment. The PCA plot was generated using *logNormCounts* and the *runPCA* function provided in the Splatter package.

### Performance Benchmarking

We conducted three compute performance benchmarks to assess the contribution of increasing number of cells, features and annotation-sets (e.g., clustering solutions) on overall run time. The impact of varying number of cells was tested using Wang et al. 2020, which has 46,500 cells and 4 different annotations on randomly sampled 10,000, 20,000, 30,000 and 40,000 cells using the scTriangulate *lazy_run* wrapper function (*assess_pruned=False, viewer_cluser=False, viewer_heterogeneity=False*). The impact of varying number of features was tested in the multiome PBMC (GEX+ATAC) using 3 annotations on the 10,991 cells, downsampled to 40,000, 60,000, 80,000 and 100,000 features and the above test function. The impact of varying number of annotation-sets was tested using the total nucleated cells CITE-Seq (donor 1), using 3,6,9 and 12 annotation-sets of varying Leiden resolutions. Runtime was evaluated using the default multi-core setting and compared with the actual CPU time if no parallelization was applied.

### Preprocessing and Quality Control

#### CITE-Seq Total Nucleated Cell

De-identified human granulocyte colony-stimulating factor (G-CSF) mobilized total nucleated cells (TNCs) were obtained through apheresis from Cell Processing Core (CPC) at Cincinnati Children’s Hospital Medical Center (CCHMC). Two young adult donors (∼30 years of age, male and female) were separately profiled. For each donor, 100,000 cells were stained with 31 human TotalSeq-A antibodies (BioLegend®, San Diego, CA) cocktail for 30 minutes on ice, and subsequently washed with flow buffer (1% FBS in PBS) and counted. ∼15,000 cells were loaded on a 10x Chromium controller with 3’ GEX version 3 chemistry. The GEX libraries were sequenced with a target depth of >50,000 reads/cell with PE150 on Illumina S4 flow cell. ADT library was sequenced with a target depth of >10,000 reads/cell with PE50 on Illumina SP flow cell.

| Antibody | BioLegend<br>TotalSeq<br>Cat # | Clone |
| --- | --- | --- |
| CD2 | 309229 | TS1/8 |
| CD3 | 300475 | UCHT1 |
| CD4 | 300563 | RPA-T4 |
| CD7 | 343123 | CD7-6B7 |
| CD8 | 301067 | RPA-T8 |
| CD10 | 312231 | HI10a |
| CD11b | 301353 | ICRF44 |
| CD13 | 301729 | WM15 |
| CD14 | 301855 | M5E2 |
| CD15 | 125615 | MC-480 |
| CD16 | 302061 | 3G8 |
| CD19 | 302259 | HIB19 |
| CD20 | 302359 | 2H7 |
| CD24 | 311137 | ML5 |
| CD33 | 366629 | P67.6 |
| CD34 | 343537 | 581 |
| CD38 | 303541 | HIT2 |
| CD45 | 304064 | HI30 |
| CD45RA | 304157 | HI100 |
| CD49f | 313633 | GoH3 |
| CD56 | 392421 | QA17A16 |
| CD64 | 305037 | 10.1 |
| CD66b | 392905 | 6/40c |
| CD71 | 334123 | CY1G4 |
| CD90 | 328135 | 5E10 |
| CD117 | 313241 | 104D2 |
| CD123 | 306037 | 6H6 |
| CD135(FLT3) | 313317 | BV10A4H2 |
| CD235a | 349117 | HI264 |
| HLADR | 307659 | L243 |
| CD110 MPL | XXXXXX | S16017A |

Raw RNA-Seq FASTQ files were aligned with Cell Ranger 3.1.0 to the human reference genome (hg19) to produce a filtered counts matrix. Cell barcode filtering, normalization, and PBMC Azimuth (level-2(L2)) reference classification were performed as with the scRNA-Seq, to yield 6,406 (replicate 1) and 4,333 (replicate 2) cells. For the ADT count, we conducted Center log-ratio (CLR) normalization using *scTriangulate*.*preprocessing*.*Normalization*.*CLR_normalization* function. We then conducted hypervariable gene selection (*top=3000*), neighbor graph construction (*n_neighbor=15*), dimension reduction (*n_pc=50*) and Leiden clustering (r=1,2,3) for RNA. In parallel, we performed neighbor graph construction (*n_neighbor=15)*, dimension reduction (*n_pc=15*) and Leiden clustering (r=1,2,3) for the ADT data. Finally, we concatenate both properly normalized RNA and ADT values to a combined AnnData through *scTriangulate*.*preprocessing*.*concat_rna_and_other* function. The combined AnnData serves as the input for the downstream scTriangulate analysis (*win_fraction_cutoff=0*.*25, add_metrics=None*). All the preprocessing and QC steps were conducted using scTriangulate version 0.9.0 and scanpy version 1.7.2. The Seurat WNN analysis was performed following the official vignette (https://satijalab.org/seurat/articles/weighted_nearest_neighbor_analysis.html). Specifically, we chose 50 Principal Components (PC)s for RNA and 18 PCs for ADT, respectively. We adjusted the resolution arguments in the FindClusters function in Seurat Version 4.0 to achieve different clustering results.

#### Single-Cell RNA-Seq of Pediatric AML CD34+ Progenitors

CD34+ positive flow cytometry sorted progenitors were isolated from whole blood from a pediatric AML patient resistant chemotherapy at the time of diagnosis. The protocol for collection and analysis was approved by the CCHMC IRB. ∼15,000 cells were loaded on a 10x Chromium controller with 5’ GEX version 1 chemistry. The GEX libraries were sequenced with a target depth of >80,000 reads/cell with PE150 on Illumina S4 flow cell (884,687,848 total reads). Raw RNA-Seq FASTQ files were aligned with Cell Ranger 2.1.1 to the human reference genome (hg19) to produce a filtered counts matrix. Distinct cell populations were identified and annotated using the software ICGS2 in AltAnalyze, using default parameters. Exon-exon junction counts for each cell barcode were obtained using the python library Pysam and imported into AltAnalyze for percent spliced-in (PSI) quantification using the MultiPath-PSI algorithm. Out of 346,972 detected exon-exon junctions associated with defined genes (Ensembl 72), 17,519 variable splicing-events with PSI estimates were reported using default options in AltAnalyze. To identify informative splicing events that are enriched in a cell population and potentially further subdivide existing clusters, we calculated pairwise comparisons for splicing events with PSI values detected in ≥ 25 cells, for each cellHarmony assigned cluster versus the largest cluster (MultiLin), first ignoring missing values (not detected in the cell) and then imputing missing values from median measurement of each event across all cells in AltAnalyze. We retained splicing events with a δPSI>0.1 and empirical Bayes t-test p<0.05, FDR corrected, with matching significant comparison prediction in both the missing-value and imputed analysis for downstream sub-clustering (n=418 events). These events were augmented with those consistently identified with the algorithm MarkerFinder using missing value PSI and imputed (n=66). To define candidate sub-clusters, we performed non-negative matrix factorization in the software ICGS2, to obtain 3 sub-clusters for each cellHarmony defined cell-population (k=3) as input for scTriangulate. For scTriangulate, we concatenate both properly normalized RNA and pre-imputed PSI values to a combined AnnData through *scTriangulate*.*preprocessing*.*concat_rna_and_other* function. The combined AnnData serves as the input for the downstream scTriangulate analysis (*win_fraction_cutoff=0*.*25*). The Seurat WNN analysis was performed following the official vignette (https://satijalab.org/seurat/articles/weighted_nearest_neighbor_analysis.html).

Specifically, we chose 50 Principal Components for RNA and 30 PCs for PSI, respectively. We adjusted the resolution arguments in the FindClusters function in Seurat Version 4.0 to achieve different clustering results.

#### Single-Cell RNA-Seq and Genotyping of CD34+ MPN Progenitors

Pre-processed count matrices for gene expression and genotyping of transcriptomes were downloaded from GEO (GSE117825). For each mutation, both wild-type and mutant allele genotypes were used to assign each cell to one of four clonal genotypes (WT, SF3B1 mutant, SF3B1-CALR mutant or SF3B1-CALR-NFE2 mutant). These mutation assignments include both false negatives (shallow sequencing) and false positives (ambient RNAs). Cell barcode filtering and normalization were performed in the software ICGS2 with default parameters and cells assigned to Hay et al. defined CD34+ cell populations using the software cellHarmony with default options. For scTriangulate, supplied clusters consist of those from cellHarmony and subclusters for cells from the assigned clonal genotypes. First, we concatenate both properly normalized RNA and genotype assignments to a combined AnnData object through the *scTriangulate*.*preprocessing*.*concat_rna_and_other* function. The combined AnnData serves as the input for the downstream scTriangulate analysis (*win_fraction_cutoff=0*.*25*).

#### scRNA-Seq PBMC

Pre-processed sparse-matrix counts (h5) were obtained from a human 10x Chromium (3’ version 3 chemistry) PBMC sample, available from the 10x Genomics website (https://support.10xgenomics.com/single-cell-gene-expression/datasets/3.0.0/pbmc_10k_v3). Cell barcodes were filtered based on a min_gene > 300, min_count > 500, pct_counts_mt < 20%, to yield 11,022 single cells. The raw UMI count matrix was adjusted to logarithmic Counts Per Ten Thousand (CPTT), we then performed hypervariable genes selection (top 3000), neighbor graph construction (*n_neighbor=15*), dimension reduction (*n_pc=50*), and Leiden clustering sequentially through the scTriangulate preprocessing module wrapper function scanpy_recipe. Separate leiden clusters (annotations) were obtained using a resolution of 1,2 and 3. All the preprocessing and Quality Control (QC) were run in scTriangulate version 0.9.0 and scanpy version 1.7.2. The AnnData after Leiden clustering was subjected to scTriangulate (*win_fraction_cutoff=0*.*25, add_metrics=‘tfidf5’*) as input. Azimuth L2 annotations were derived from the filtered raw count data through the Azimuth PBMC online R-Shiny app (https://app.azimuth.hubmapconsortium.org/app/human-pbmc).

#### Multiome PBMC

Multiome PBMC sparse matrix counts h5 data was downloaded from the 10x Genomics website (https://www.10xgenomics.com/resources/datasets/pbmc-from-a-healthy-donor-granulocytes-removed-through-cell-sorting-10-k-1-standard-2-0-0). We conducted QC based on both RNA and ATAC peaks. We filtered out nuclei with min_genes < 300, min_counts < 500, pct_counts_mt > 20% for RNA data, together with the additional criteria for at least 1000 peaks/nucleus in the ATAC data based on episcanpy ^38^ tutorial. Taken together, 10,991 nuclei were kept for further analysis. We normalized both RNA and ATAC count data to logarithmic CPTT, then performed hypervariable gene selection (top=3000), neighbor graph construction (*n_neighbor=15*), dimension reduction (*n_pc=15*), and Leiden clustering (r=1,2,3) for the RNA expression. For the ATAC peaks, we performed hypervariable peak selection (top=100,000), neighbor graph construction (*n_neighbor=15*), dimension reduction (n_pc=100) and Leiden clustering (r=1,2,3). The arguments for n_top_features and n_pc was performed as indicated in the episcanpy^38^ tutorial. Finally, we concatenate both properly processed RNA and ATAC values to a combined AnnData through *scTriangulate*.*preprocessing*.*concat_rna_and_other* function. The combined AnnData serves as the input for the downstream scTriangulate analysis (*win_fraction_cutoff=0*.*35, add_metrics=‘tfidf5’*). The inclusion of the secondary TF-IDF score was based on a preliminary evaluation with both the default and additional tfidf5 parameters, which better reflected known cell population marker diversity. All the preprocessing and QC steps were conducted using scTriangulate version 0.9.0 and scanpy version 1.7.2.

#### TEA-Seq PBMC

TEA-Seq provides simultaneous measurement of RNA, ADT, and ATAC in the same single nuclei. The RNA-ATAC combined h5 files and ADT count CSV files were downloaded from NCBI GEO (GSM4949911). Customized scripts (available on Github) were used to find the common nuclei in these two files and three different AnnData corresponding to three modalities were generated. We conducted QC for both RNA and ATAC, based on the same cutoffs mentioned above, nuclei with *min_gene* < 300, *min_counts* < 500, *pct_counts_mt* > 20%, *min_atac_peaks* < 1000 will be removed. This step had 8,213 nuclei retained for further analysis. For the RNA, we normalized the RNA UMI count to logarithmic CPTT, we then performed hypervariable gene selection (*top=3000*), neighbor graph construction (*n_neighbor=15*), dimension reduction (*n_pc=15*), and Leiden clustering (r=1,2,3). For the ATAC, we first normalized the ATAC peak count to logarithmic CPTT, we then performed hypervariable peak selection (top=60000), neighbor graph construction (*n_neighbor=15*), dimension reduction (*n_pc=100*), and Leiden clustering (r=1,2,3). The arguments for n_top_features and n_pc base on both the episcanpy ^38^ tutorial and dedicated experiments. For the ADT, we conducted CLR normalization using the *scTriangulate*.*preprocessing*.*Normalization*.*CLR_normalization* function, followed by neighbor graph construction (*n_neighbor=15*), dimension reduction (*n_pc=15*) and Leiden clustering (r=1,2,3). Finally, we combine three processed AnnData objects using scTriangulate.preprocessing.concat_rna_and_other function. The combined AnnData serves as the input for the downstream scTriangulate analysis (*win_fraction_cutoff=0*.*25, add_metrics=None*). The Azimuth L2 annotation was obtained using post-QC RNA raw UMI count data through the web app (https://app.azimuth.hubmapconsortium.org/app/human-pbmc). All the preprocessing and QC steps were conducted using scTriangulate version 0.9.0 and scanpy version 1.7.2. The Seurat 3-way WNN analysis (RNA, ADT, ATAC) was performed following the official vignette (https://satijalab.org/seurat/articles/weighted_nearest_neighbor_analysis.html). Specifically, we chose 50 Principal Components (PC)s for RNA, 18 PCs for ADT and 2-50 dimensions of latent spaces (Latent semantic analysis) for ATAC, respectively. We adjusted the resolution arguments in the FindClusters function in Seurat Version 4.0 to obtain different clustering results. The totalVI analysis was performed on only RNA and ADT modalities following the official tutorial (https://docs.scvi-tools.org/en/stable/tutorials/notebooks/totalVI.html) using scvi_tools version 0.14.3. Specifically, we chose the top 4000 variable genes as the RNA features and 400 training epochs (default). We performed the Leiden clustering on the totalVI learned joint embedding space and adjusted the resolution arguments in *scanpy*.*tl*.*leiden* function to obtain results for a range of clustering resolutions. For evaluation of scTriangulate relative to popular ensemble-clustering algorithms, all 9 independent Leiden clustering results derived from the three independent TEA-Seq modalities (RNA, ATAC, ADT) were provided to the HyperGraph Partitioning Algorithm (HGPA), Meta-Clustering Algorithm (MCLA) and Hybrid Bipartite Graph Formulation (HBGF), using the Python package ClusterEnsembles with default options (https://github.com/tsano430/ClusterEnsembles). Cluster-based similarity Partitioning Algorithm (CSPA) was excluded through the evaluation due to its poor suitability for large datasets.

#### Lung Cell Atlas

scRNA-Seq and snRNA-Seq cell-population assignment and counts data were obtained from four independent studies (GSE161383, GSE171524, GSE136831, EGAS00001004344). Among these, a lung snRNA-Seq from 9 donors of varying age (newborn, ∼3 years old, and ∼30 years old)^25^ was selected as the base dataset, whereas the remaining were classified on to this query dataset using the software cellHarmony. In brief, counts for each dataset were scaled to CPTT and log2 adjusted. For each dataset, reference centroids were initially computed from the author-provided cell-type labels by identifying the top-60 marker genes (MarkerFinder algorithm) for non-diseased control samples (excluding cell-cycle gene enriched clusters). cellHarmony was run in two steps: 1) centroid-based classification of each reference to Wang et al. 2020 with a Pearson correlation rho>0.2 and 2) self-reclassification of Wang et al. 2020 to the re-derived markers and centroids from the first classification (MarkerFinder and cellHarmony), to improve the accuracy of the classifications between scRNA-Seq and snRNA-Seq. The same two-step mapping strategy was applied to HLCA (https://app.azimuth.hubmapconsortium.org/app/human-lung-v2), LungMAP CellRef 1.0 (https://app.lungmap.net/app/shinycell-human-lung-cellref) primary Azimuth mappings. Similarly, to assess the quality of scTriangulate myeloid cell population annotations, we compared these specifically to a curated set of Mononuclear phagocytes (MNP-Verse) ^30^. Here, we computed gene expression centroids from only healthy lung control samples provided by Mulder et al. and mapped the Wang et al. snRNA-Seq dataset using this two-step approach ^39^. The obtained annotations and UMAP coordinates ^25^ were added to the existing AnnData by the *scTriangulate*.*preprocessing*.*add_annotation* and *scTriangulate*.*preprocessing*.*add_umap* functions respectively. The RNA raw UMI count was normalized to logarithmic CPTT. The processed AnnData was used as the input for running scTriangulate (*win_fraction_cutoff=0*.*4, add_metrics=‘tfidf5’*). The inclusion of the secondary TF-IDF score was based on a preliminary evaluation with both the default and additional tfidf5 parameters, which better reflected known cell population marker diversity.

#### Bone Marrow Cell Atlas

Alternative scRNA-Seq clustering algorithm results and curated annotations were obtained from our prior benchmarking study ^8^, using results deposited in the Synapse database (syn26320732). The obtained annotations and umap coordinates were added to the existing AnnData by scTriangulate.preprocessing.add_annotation and *scTriangulate*.*preprocessing*.*add_umap* functions respectively. The RNA raw UMI count was normalized to logarithmic CPTT. The processed AnnData was used as the input for running scTriangulate (*win_fraction_cutoff=0*.*3, add_metrics=‘tfidf5’*). Agreement of scTriangulate using results from ICGS2, Monocle3 and Seurat3 with Hay et al. clusters was performed using default parameters (**Supplementary Figure 4d**).

---

**Supplementary Figure 1.**
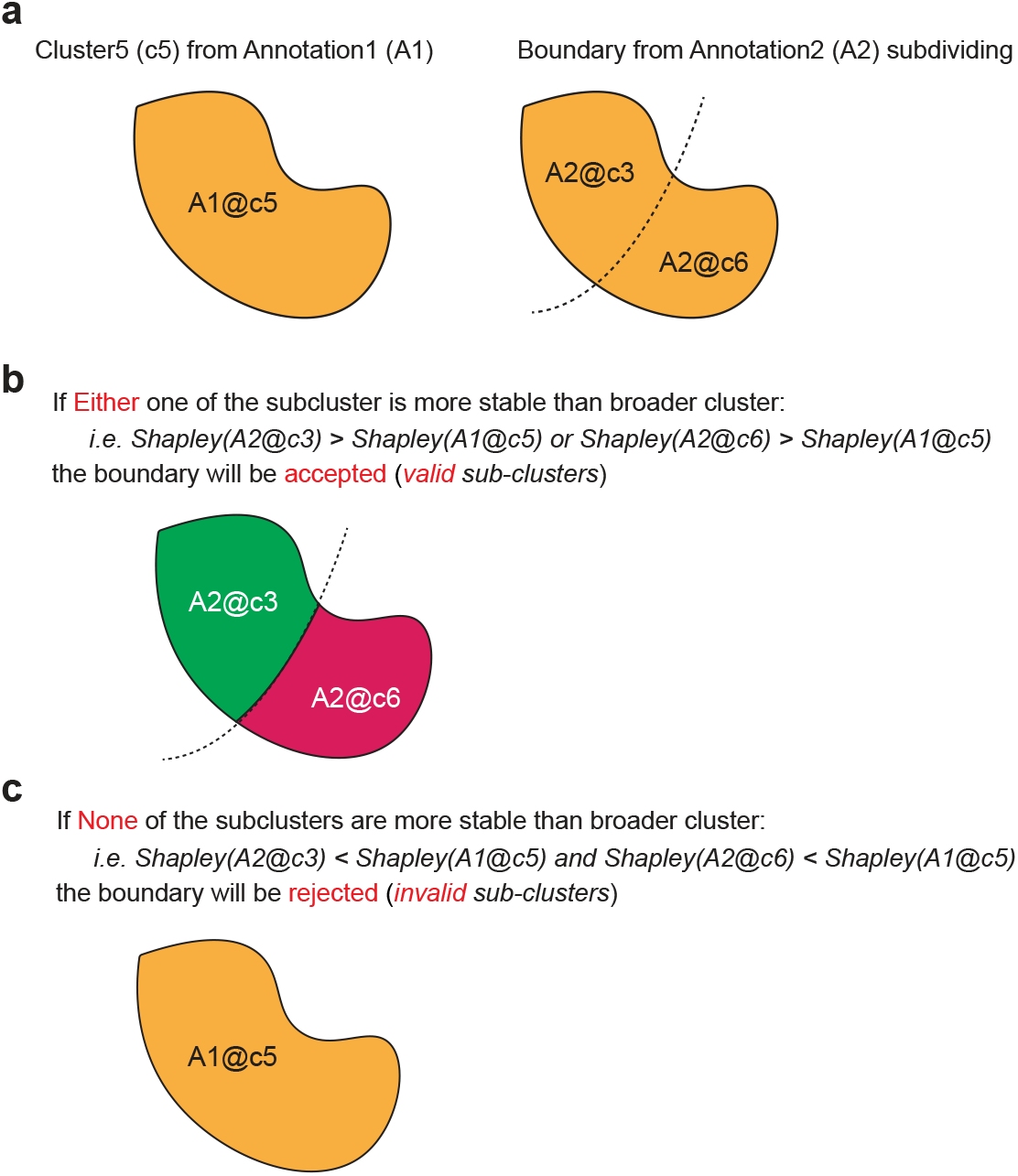
Defining stable versus unstable subclusters with scTriangulate. A theoretical example is outlined for the determination of stable subclusters via comparison of the Shapley Value statistic. a) One cluster (c5) in annotation-set A1 (left) is divided by an imaginary boundary from an overlapping annotation-set (A2) (right). b) Under the condition where either one of the subclusters (A2@c3, or A2@c6) is more stable than the broader cluster (A1@c5), based on the Shapley Value derived from the reassign, TF-IDF, and SCCAF scores, this boundary will be kept and serve as a valid division that demarcates two stable subpopulations. c) Alternatively, under the condition where none of the subclusters is more stable than the broader cluster, the program will reject this boundary.

**Supplementary Figure 2.**
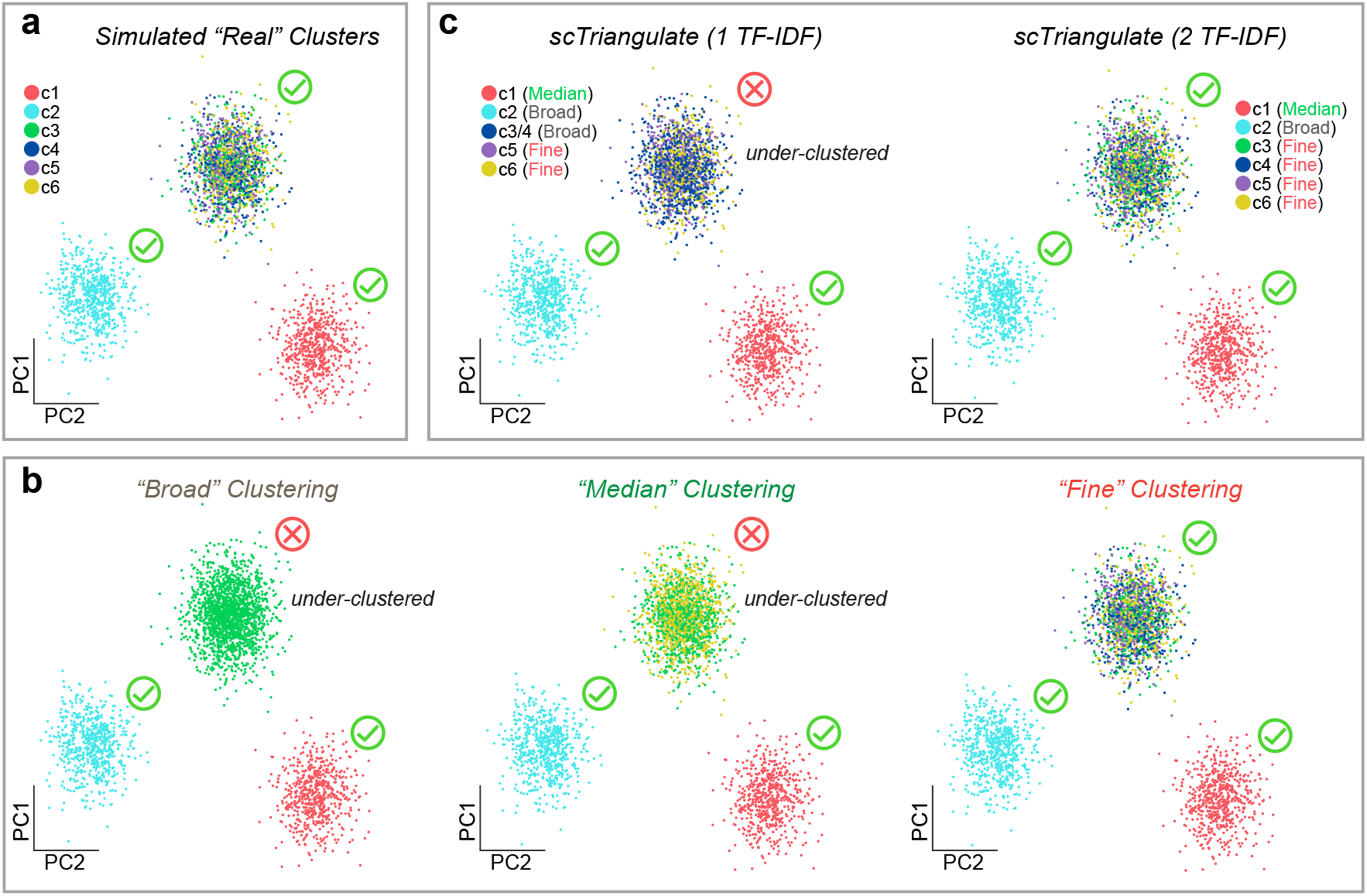
Benefit of additional stability metrics in the game. a) As a ground-state truth to assess the specificity of predictions by scTriangulate, we simulated scRNA-Seq data for two highly distinct simulated populations (c1 and c2) and four subtly different subclusters (c3, c4, c5, c6) (Methods). b) To predict such clusters, we produced three cluster annotations for the same cells in panel a, with different granular cluster definitions (broad, median and fine). c) scTriangulate integration of Broad, Median and Fine clusters, using the default two TF-IDF scores (TF-IDF10 and TF-IDF5) or one TF-IDF score alone (TF-IDF10).

**Supplementary Figure 3.**
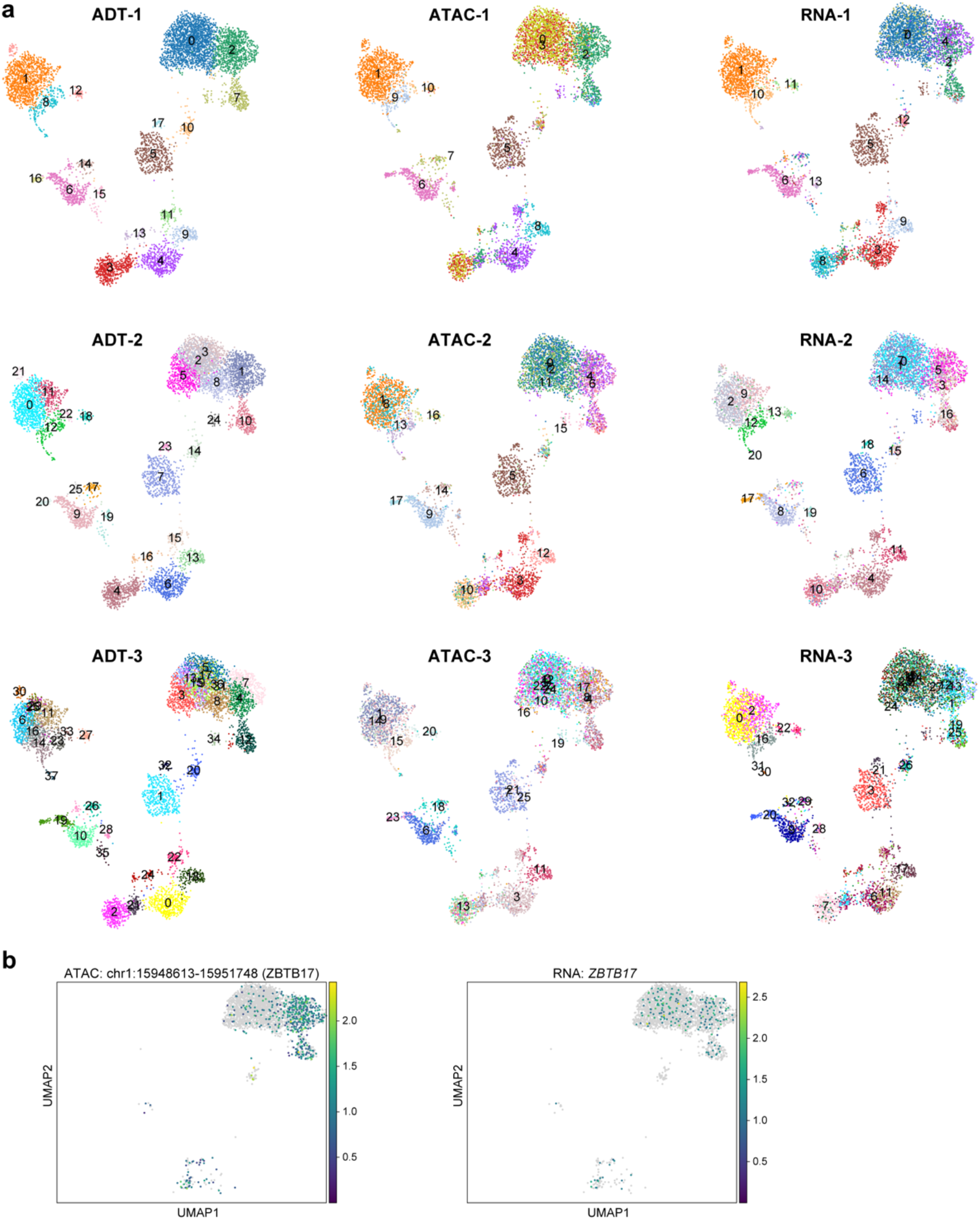
Trimodal integration of PBMC clusters. a) Scanpy Leiden UMAP for multimodal analysis of human blood by TEA-Seq analysis (RNA+ATAC+ADT). UMAP coordinates are derived from the ADT data. Clustering results from three RNA, three ATAC and three ADT resolutions (r=1,2,3) are shown. b) Visualization of the top multiome ATAC marker (critical early T cell progenitor regulatory factor ZBTB17 ^18^) for the CD95+ subset (left) and corresponding *ZBTB17* gene expression in the same genomic loci and nuclei.

**Supplementary Figure 4.**
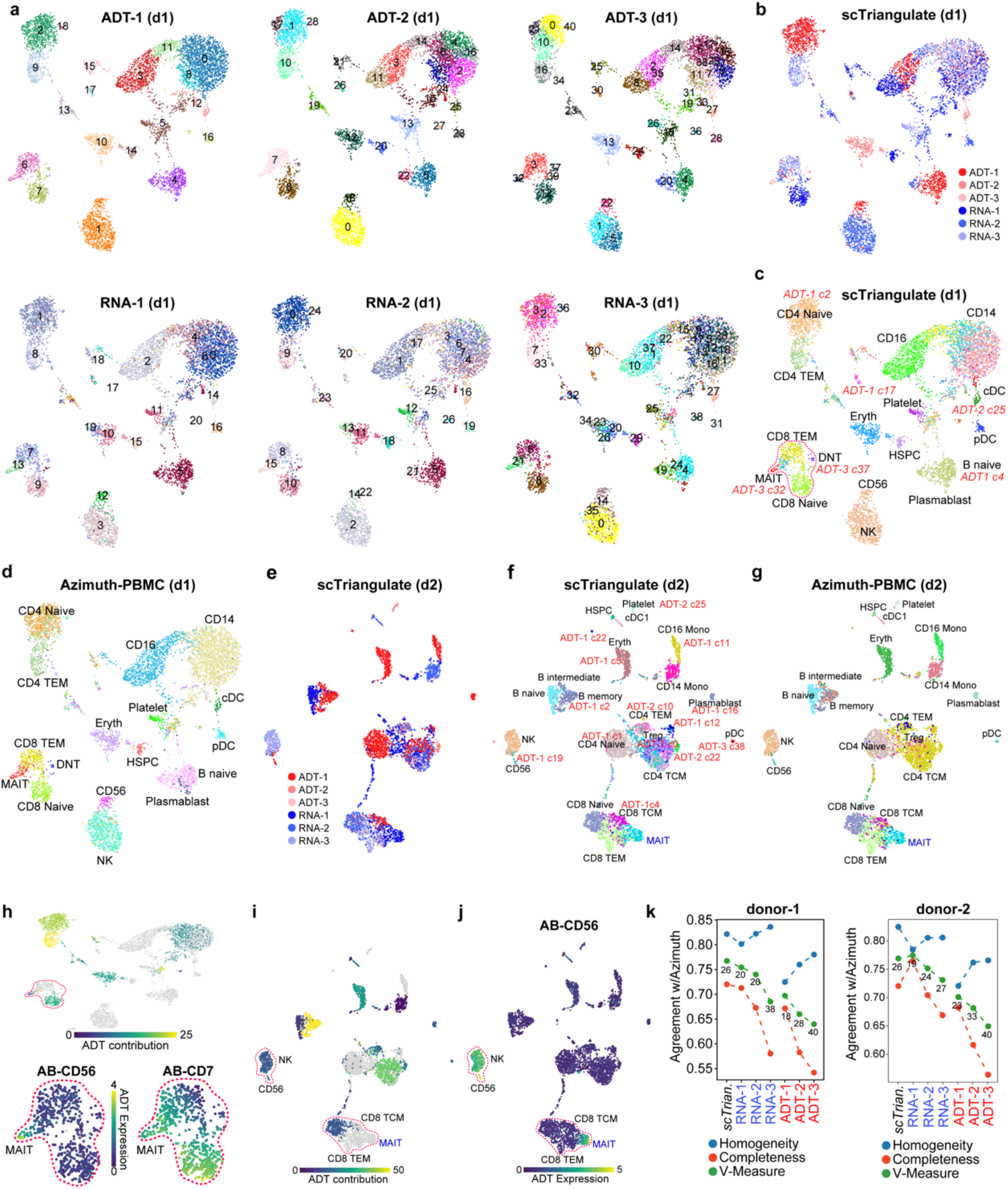
Novel lymphoid subsets resolved through multimodal CITE-Seq integration. a) UMAP of CITE-Seq total nuclear cells (TNC) from donor 1, produced by scanpy Leiden clustering of three RNA and three ADT resolutions (r=1,2,3). b) scTriangulate final clusters with labels colored according to the source Leiden cluster and modality (donor 1). c) scTriangulate winning clusters after integration of multiple clustering resolutions (n=3) for each assayed modality relative to Azimuth annotations (donor 1). Clusters specifically derived from ADT cluster resolutions are labeled in the plot. d) Reference cell population labels from PBMC level-2 Azimuth projection (donor 1). e-g) Same as panels b,c and d, respectively, for CITE-Seq of donor 2. h) The selective contribution of ADTs is overlaid on the CITE-Seq UMAP (donor 1), based on the frequency of associated features among the top-20 markers of each final cluster (Methods). In the lower panel, the specificity of specific ADTs (CD56, CD7), demarcates the cluster boundaries of scTriangulate-defined MAIT cell subpopulations and CD8 T Effector Memory cell subpopulations, respectively. i) Relative contribution of ADT features to scTriangulate final clusters (donor 2). j) Independent evaluation of MAIT and NK CD56+ bright cell heterogeneity based on CD56 ADT expression (donor 2). k) Agreement with Azimuth PBMC reference label assignments measured by Homogeneity, Completeness, and V-Measure, for each individual or integrated set of clustering solutions (donor 1 left, donor 2 right). The number of clusters produced by each indicated resolution are displayed above the V-Measure data point.

**Supplementary Figure 5.**
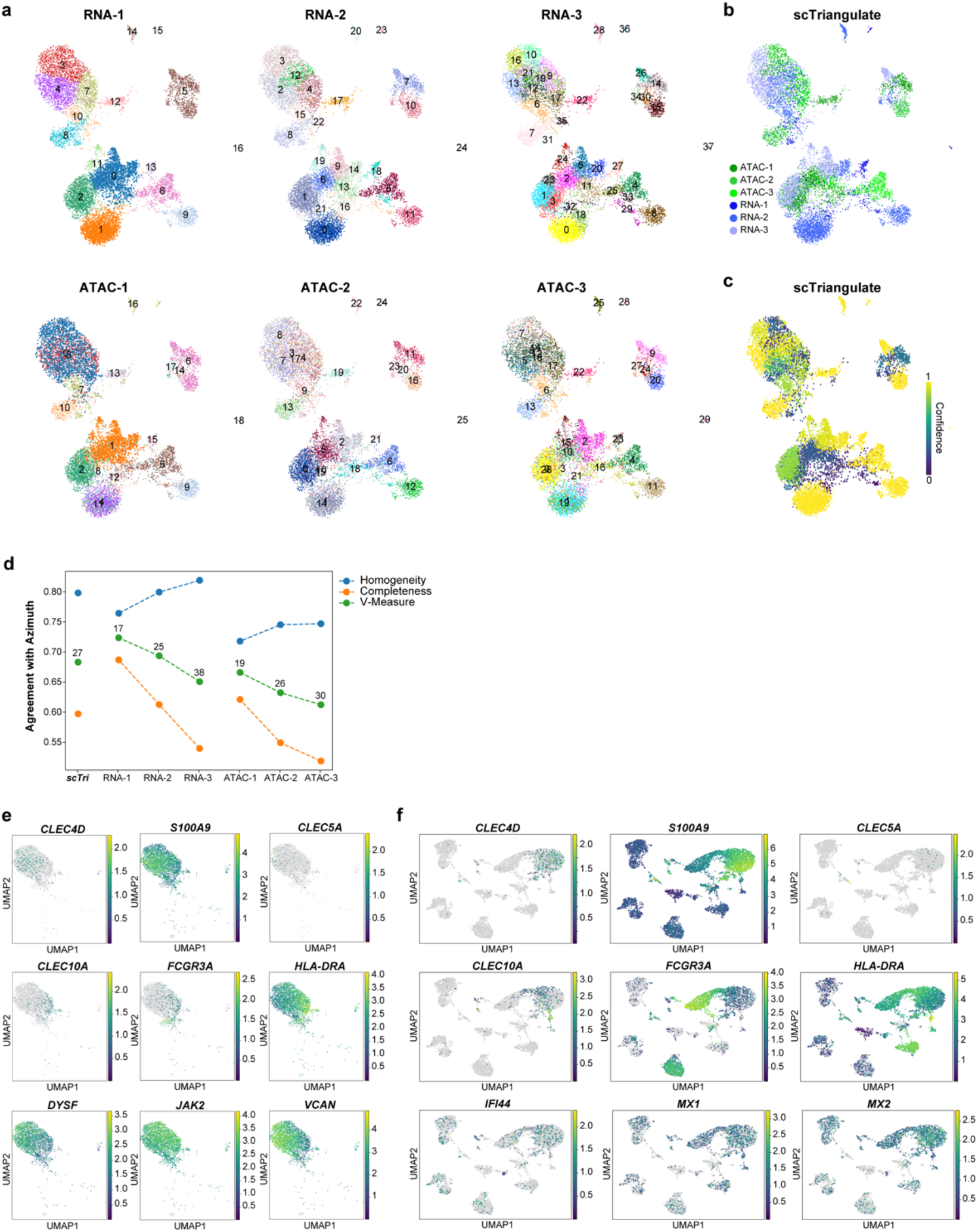
New CD14 monocyte subsets resolved through the integration of ATAC- and RNA. a) Scanpy Leiden UMAP for multimodal analysis of human blood by single-nuclei multiome analysis from three RNA and three ATAC resolutions (r=1,2,3). UMAP coordinates are derived from RNA. scTriangulate final clusters with labels colored according to the source Leiden cluster and modality. c) Cluster confidence for each final cluster (winning fraction, see Methods). d) Agreement with Azimuth PBMC reference label assignments measured by Homogeneity, Completeness, and V-Measure, for each individual or integrated set of multiome clustering solutions. e, f) Evidence for commonly detected subsets of monocytes in (e) multiome and (f) CITE-Seq with CD14 Monocytes. Canonical markers of monocyte heterogeneity with Azimuth defined CD14 Monocytes; classical CD14 Monocyte = *S100A9, CLEC4D, CLEC5A*; intermediate CD14 Monocyte = *CLEC10A, FCGR3A, HLA-DRA*; inflammatory monocyte = *MX1, MX2, IFI44* and additional scTriangulate predicted top Monocyte population markers = *DYSF, JAK2, VCAN*.

**Supplementary Figure 6.**
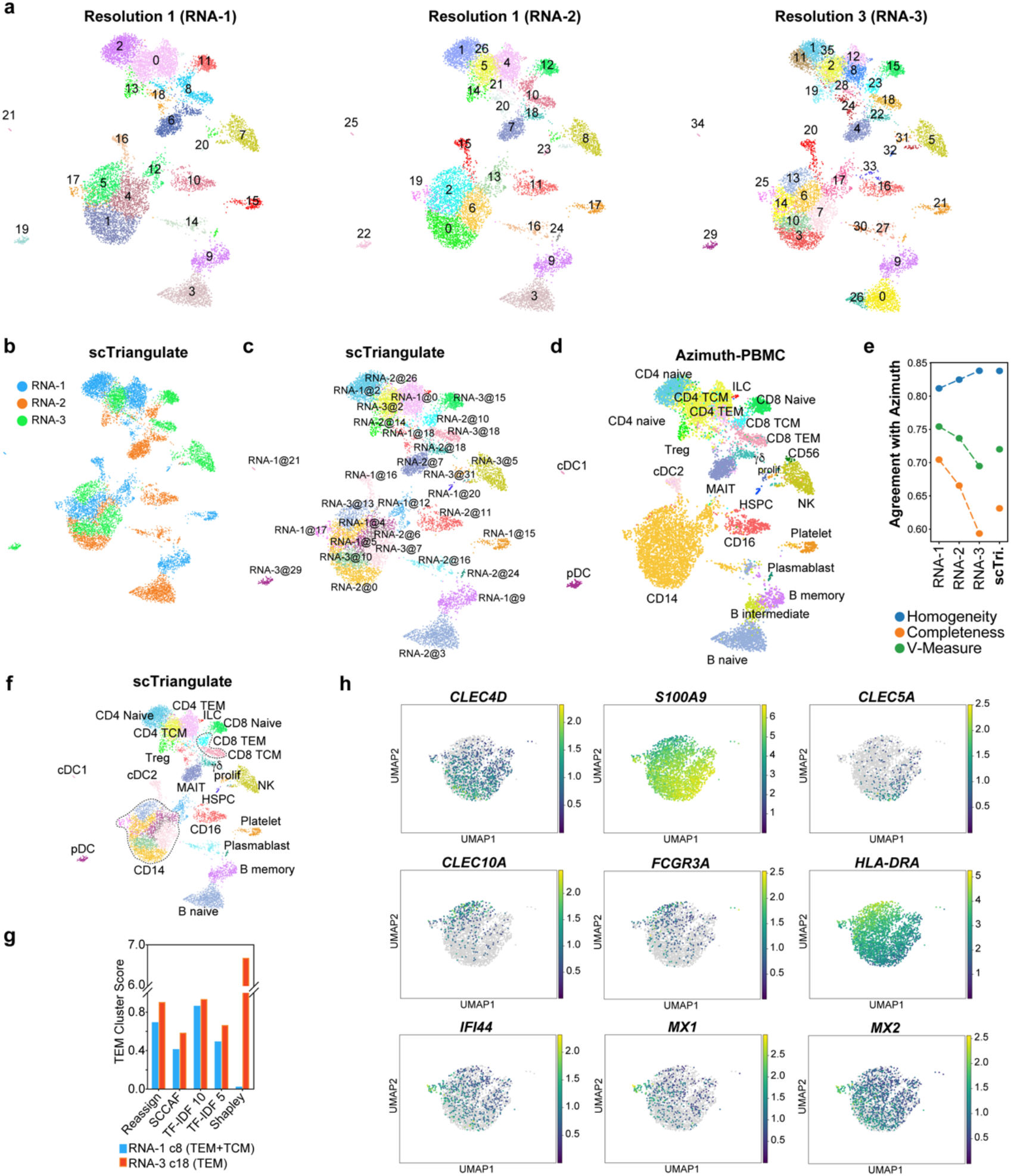
Accurate prediction of PBMC scRNA-Seq consensus populations from different software resolutions. a) Scanpy Leiden clustering results for three resolutions (RNA-1, RNA-2, and RNA-3). b) scTriangulate final cluster source considering all evaluated resolutions. c) scTriangulate defined clusters after pruning and reclassification. The prefix of each label indicates the annotation source and the suffix the original cluster label from panel a. d) Projected labels from a PBMC CITE-Seq reference (Azimuth, Level 2). e) Agreement with Azimuth PBMC reference label assignments measured by Homogeneity, Completeness, and V-Measure, for each individual clustering solution. f) Overlaid cell-type annotations from Azimuth on scTriangulate results, with the delineation of rare CD8 T-cell subtypes highlighted in g and CD14 monocytes in h. g) Comparison of each stability metric and Shapley between a broader T cell memory cluster (light blue) and a more granular subset (T effector memory or TEM, red). h) Canonical markers of monocyte heterogeneity with Azimuth defined CD14 Monocytes; classical CD14 Monocyte = *S100A9, CLEC4D, CLEC5A*; intermediate CD14 Monocyte = *CLEC10A, FCGR3A, HLA-DRA*; and inflammatory monocyte = *MX1, MX2, IFI44*.

**Supplementary Figure 7.**
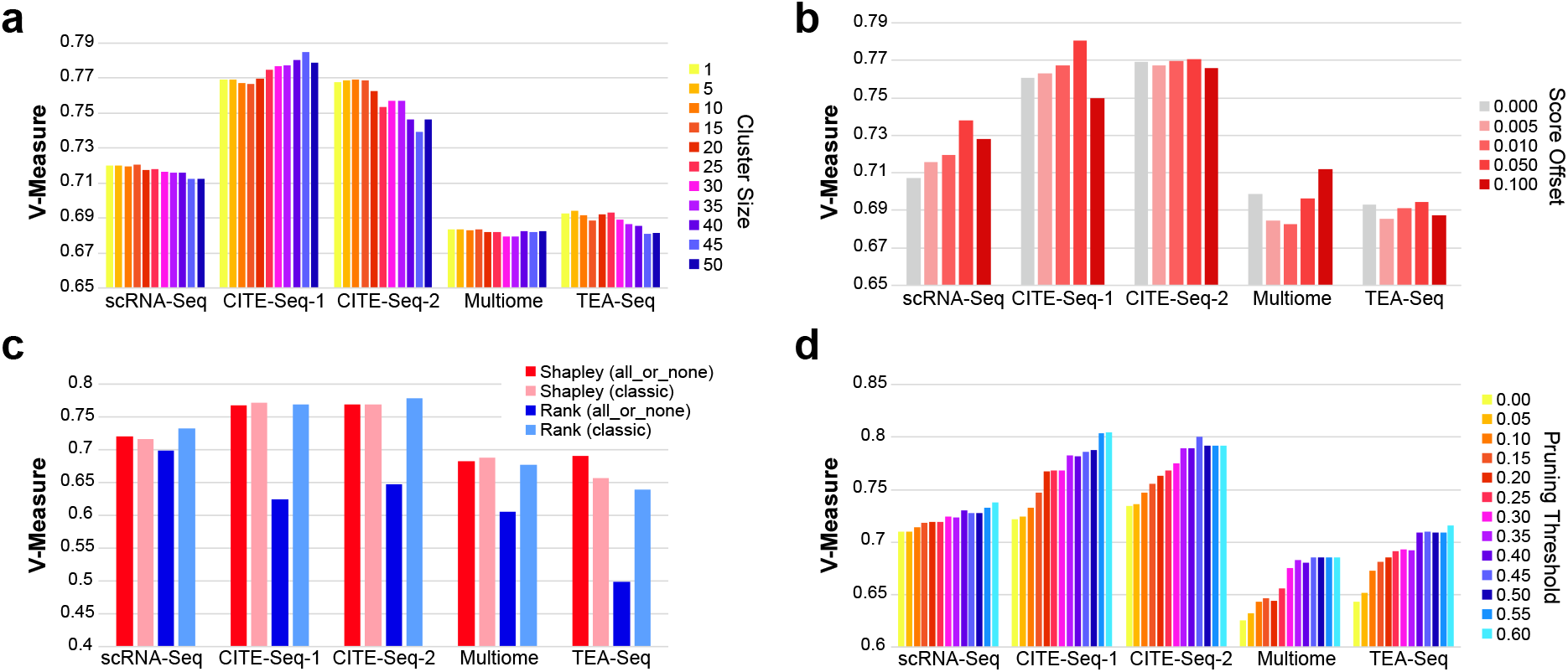
Sensitivity analysis for distinct tunable parameters in scTriangulate. The performance of different scTriangulate tunable parameters were tested for the blood single-cell genomics datasets evaluated in this manuscript (unimodal and multimodal) (Methods). Each bar indicates the correspondence of clusters in scTriangulate when compared to the PBMC Azimuth reference (quantified by V-measure). a) Comparison of cutoffs for reliable minimum number of cells in a source cluster (Cluster Size) for all evaluated annotations, ranging from 1-50 cells. b) Comparison of different stability metric rank tolerance (offset) thresholds, ranging from 0.0-0.1. c) Comparison of different annotation importance strategies (Shapley and simple Rank-based), for different rank prioritization approaches (all_or_none, classical). All_or_none = winner takes all strategy. Classic = unadjusted rank strategy. d) Comparison different cluster pruning thresholds, ranging from 0-60% of cells retained from the original parent cluster.

**Supplementary Figure 8.**
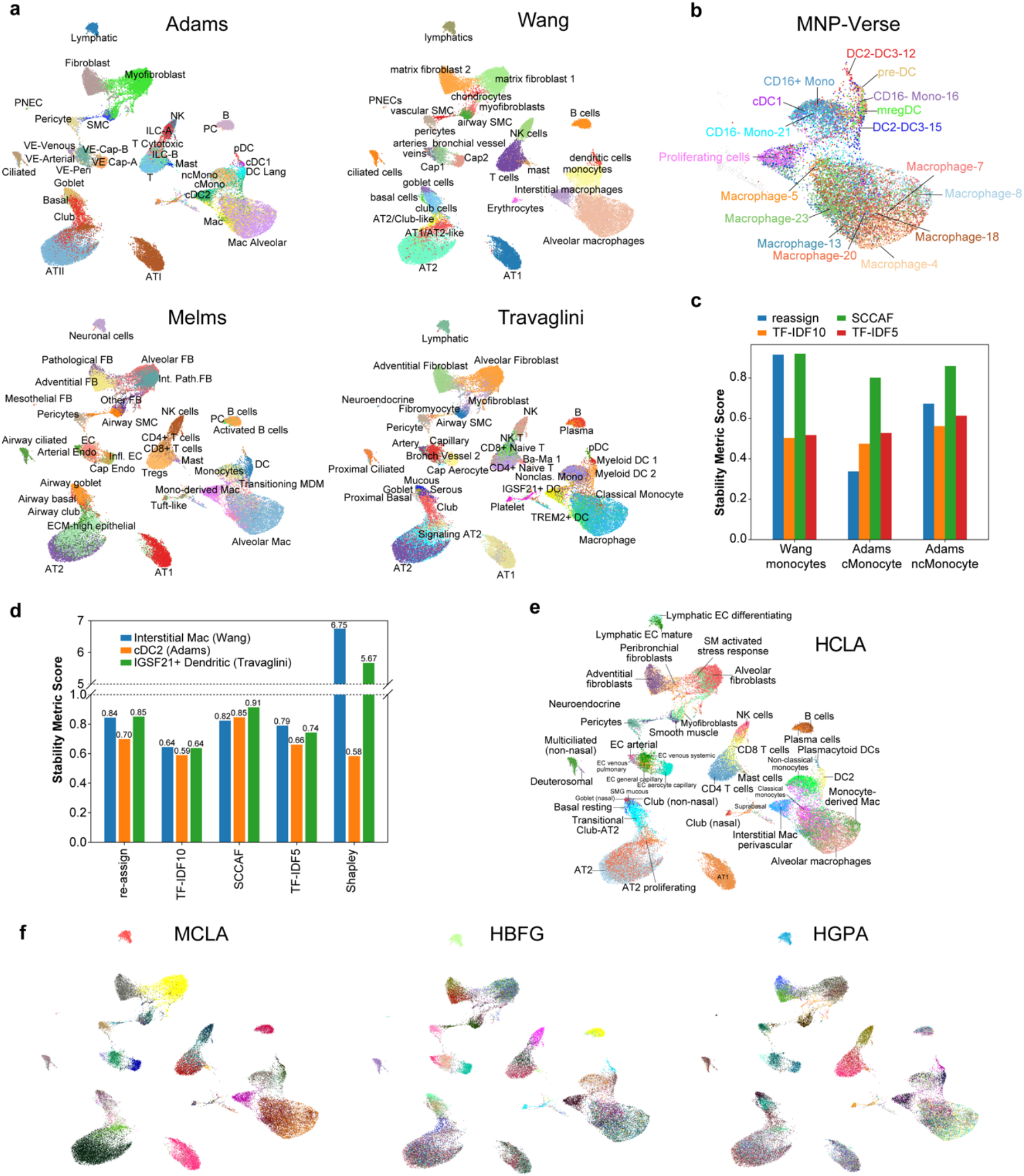
Integrating multiple lung cell annotations to create a unified cell atlas. a) cellHarmony supervised mapping of scRNA-Seq reference labels (Adams et al. 2020, Melms et al. 2021, Travaglini et al. 2020) on to Wang et al. 2020 lung snRNA-Seq. b) Cell atlas annotations from MNP-Verse (myeloid-specific reference) projected onto Wang snRNA-Seq myeloid cell populations. c) Comparison of scTrianguate stability metrics for Wang monocytes, relative to Adams et al. cMono and nMono. d) scTriangulate considered stability metrics for Wang Interstitial macrophages, compared specifically to overlapping Adams and Travaglini aligned cluster definitions. e) Cell annotations from the human lung cell atlas (HCLA) reference (Azimuth projected - finest level) onto the Wang snRNA-Seq data. f) Ensemble clustering of the supervised mapping assignments from panel a, using MCLA, HBGF and HGPA.

**Supplementary Figure 9.**
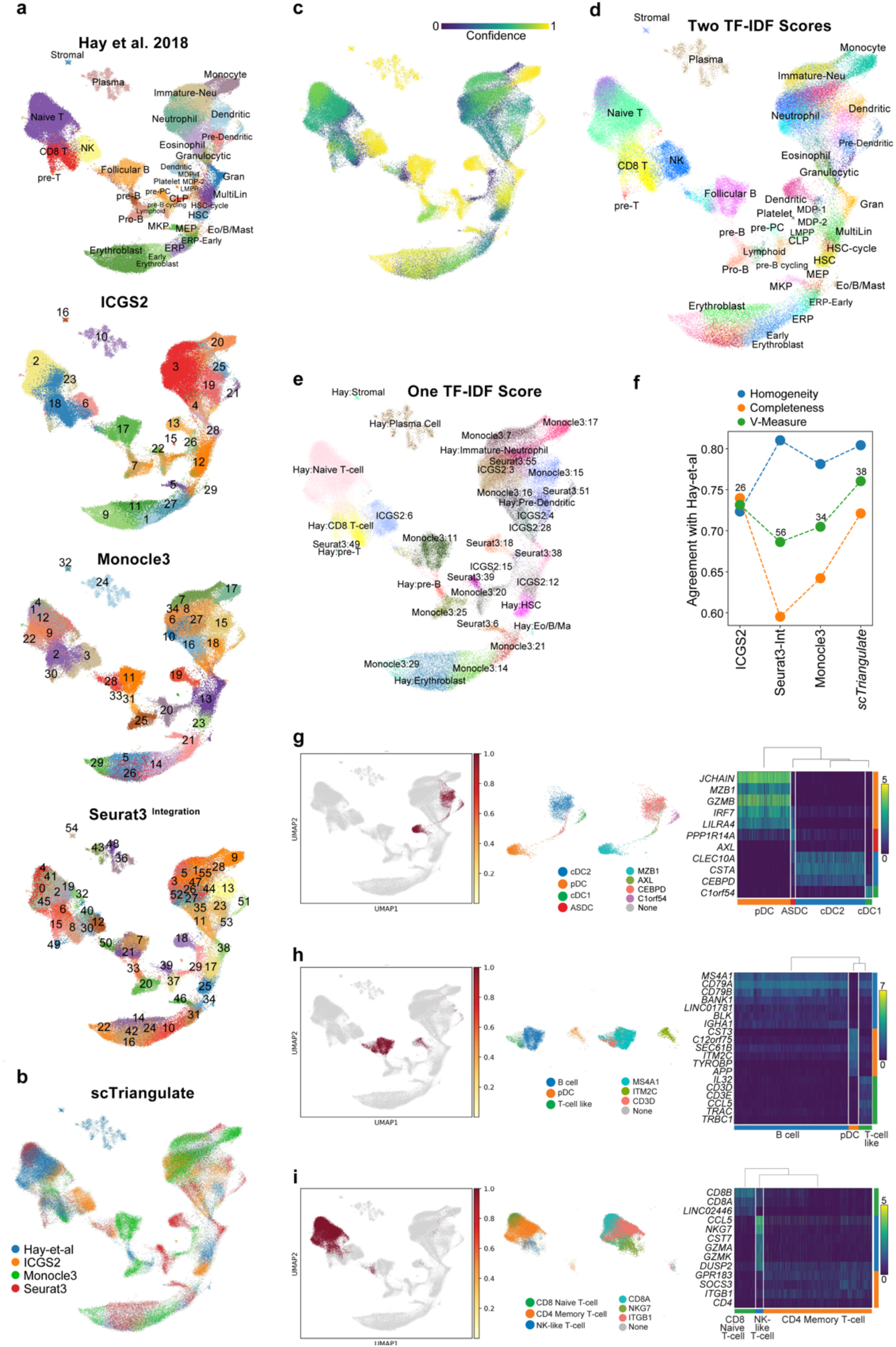
Integration of diverse single-cell algorithms and curated annotations in Bone Marrow. a) UMAP of previously produced scRNA-Seq clustering results from three algorithms (ICGS2, Seurat3 and Monocle3) with the author source annotations (Hay et al. 2018) for an adult bone marrow atlas spanning >100,000 cells, 8 donors and 64 captures (10x Chromium 3’ version 2). b) scTriangulate final cluster source from panel a. c) Cluster confidence for each final cluster (winning fraction, see Methods). d) scTriangulate results for the default setting parameters (two TF-IDF scores, TFIDF5 and TFIDF10), which are expected to produce more granular clusters (marker gene-driven Shapley Value). e) scTriangulate final results when only one TF-IDF score (TFIDF10) was applied (conservative). f) Agreement with Hay et al. cluster annotations for scTriangulate compared to Seurat 3 integration, Monocle 3 and ICGS 2, measured by Homogeneity, Completeness, and V-Measure. g-i) The original author-identified (g) Dendritic Cell (DC) cluster, (h) Follicular B cells, and (i) Naive T cells (left), split into four subpopulations (middle) with support by canonical marker-gene expression (middle).

**Supplementary Figure 10.**
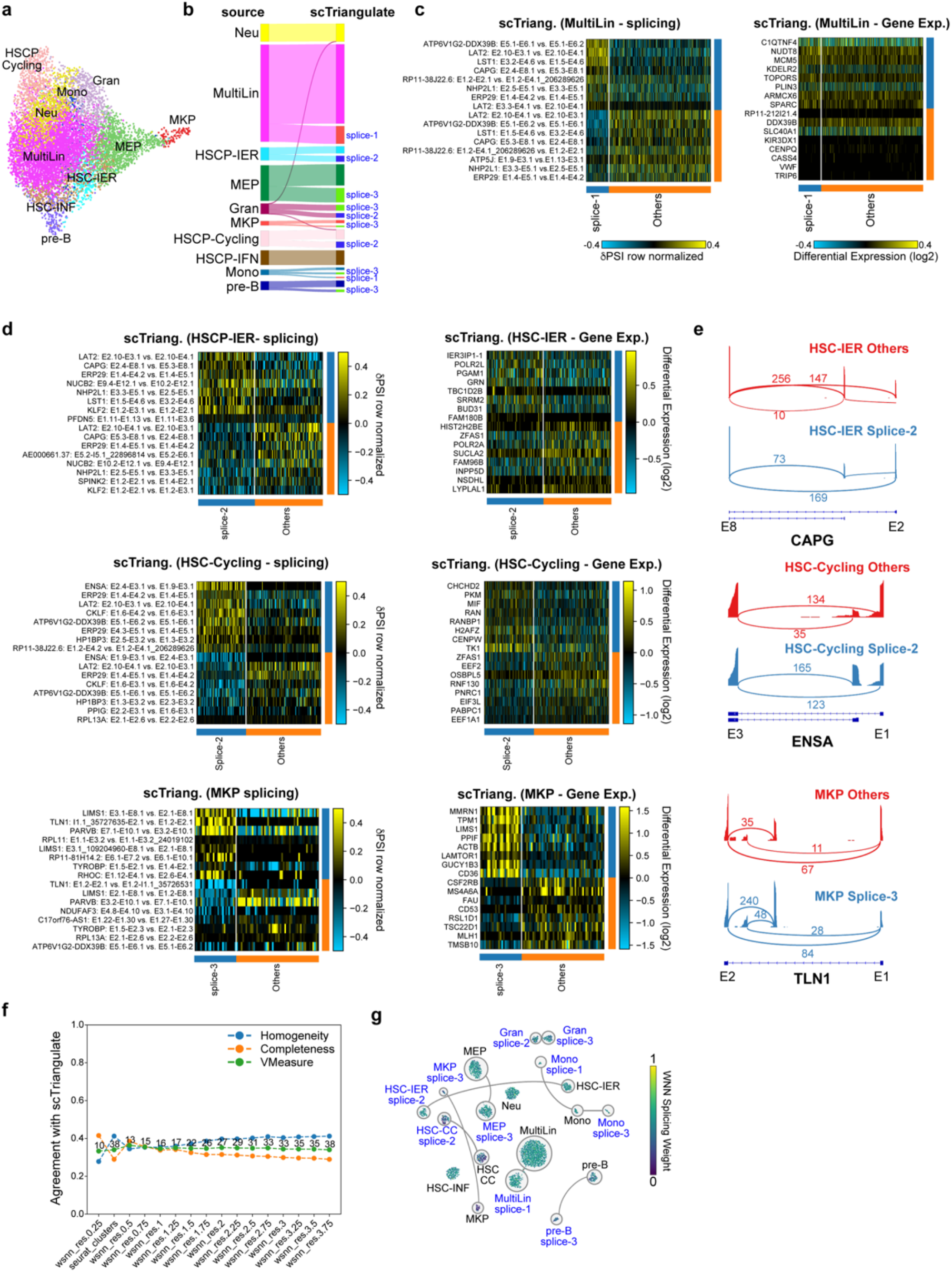
Stable-splicing defined leukemic cell-population subtypes. a) UMAP of pediatric AML scRNA-Seq, produced from cellHarmony supervised cell assignments against normal CD34+ progenitor clusters. b) Sankey diagram of scTriangulate winning-cell populations from gene expression and splicing. Splicing-defined subclusters are denoted in blue along with the source splicing-NMF cluster number (1,2 or 3). c,d) Heatmaps displaying top discriminating splicing (left) and gene expression (right) markers that subdivide cellHarmony gene-expression defined cell populations. e) Representative SashimiPlots for marker splicing-events for scTriangulate winning clusters. f) Agreement with scTriangulate cluster annotations for a range of Seurat 3 WNN integration resolutions, measured by Homogeneity, Completeness, and V-Measure. The number of clusters produced by each indicated resolution are displayed above the V-Measure data point. g) Visualization of the Seurat WNN splicing “weight” on the scTriangulate AML UMAP.

**Supplementary Figure 11.**
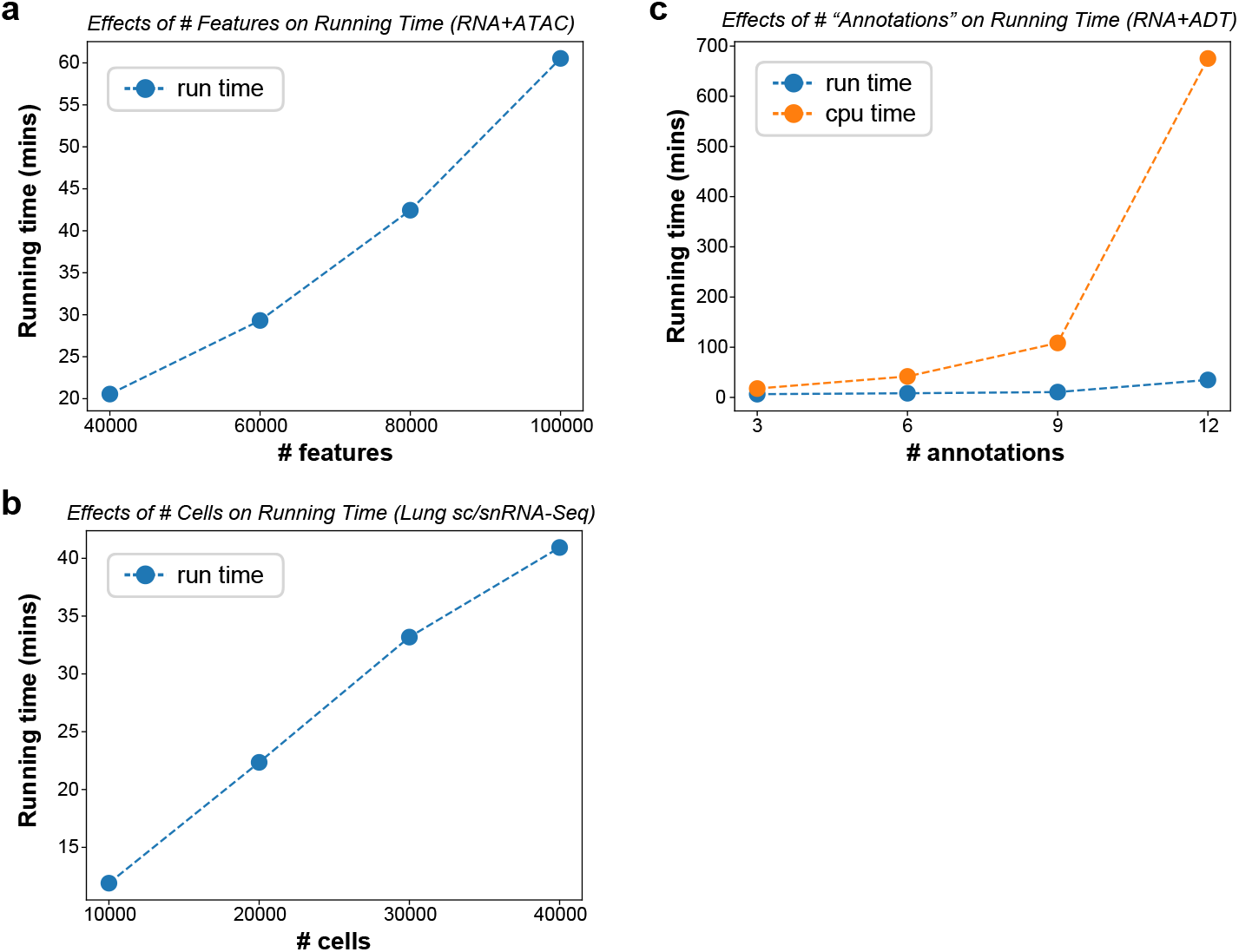
Run time and scalability of scTriangulate. a) The impact of increasing numbers of features on runtime is shown for the multiome GEX+ATAC dataset (>100,000 features) in this study with randomly downsampling. b) The impact of increasing numbers of cells on runtime is shown for the Wang et al. Lung dataset (∼46,000 single cells) with all four annotation sources, with random downsampling of cell barcodes. c) The impact of increasing numbers of annotation-sets on runtime is shown for the CITE-Seq (sample 1) dataset (∼6,000 single-cells), for 3, 6, 9, 12 query annotation-sets (in parallel, by default, blue line), compared with CPU time (sequentially, orange line).

